# Loss of polarity regulators initiates gasdermin E mediated pyroptosis in human maternal fetal interface trophoblasts

**DOI:** 10.1101/2022.06.30.498172

**Authors:** Khushali Patel, Jasmine Nguyen, Sumaiyah Shaha, Amy Brightwell, Wendy Duan, Ashley Zubkowski, Meghan Riddell

## Abstract

The syncytiotrophoblast is the placental epithelial cell that forms the maternal surface of the human placenta, acting as a barrier and facilitating exchange between mother and fetus. Syncytiotrophoblast dysfunction is a feature of pregnancy pathologies, like preeclampsia. Dysfunctional syncytiotrophoblast display a loss of microvilli, a marker of aberrant apical-basal polarization, but little data exists about the regulation of syncytiotrophoblast polarity. Atypical protein kinase-c (aPKC) isoforms are conserved polarity regulators. Thus, we hypothesized that aPKC isoforms regulate syncytiotrophoblast polarity. Using human placental explant culture and primary trophoblasts, we found that loss of aPKC activity or expression induces syncytiotrophoblast gasdermin E dependent pyroptosis. We also establish that TNF-α induces an isoform specific decrease in aPKC expression and gasdermin E dependent pyroptosis. Therefore, aPKCs are homeostatic regulators of syncytiotrophoblast function and a pathogenically relevant pro-inflammatory signal leads to a highly pro-inflammatory form of cell death at the maternal-fetal interface. Therefore, our results have important implications for the pathobiology of placental disorders.

## Introduction

The placenta is a fetally-derived transient organ that performs critical functions like gas and nutrient exchange, pregnancy supporting hormone secretion, and acts as a physical barrier between the maternal and fetal compartments to establish and maintain a healthy pregnancy.^1^ The maternal-facing surface of the human placenta is covered by a single giant multinucleated epithelial cell called the syncytiotrophoblast (ST), which measures up to 10m^2^ by the end of gestation.^2^ The ST is a highly polarized epithelial cell with dense apical microvilli decorating its surface, but it lacks lateral membranous barriers between nuclei. Thus, the ST represents a completely unique type of epithelial barrier. Since the ST cannot undergo mitosis, it is maintained by the fusion of underlying proliferative-mononuclear progenitor cytotrophoblasts. The ST is the primary cell-type responsible for carrying out the essential placental functions listed above, therefore ST stress and dysfunction is a key feature of common pregnancy complications like preeclampsia and intrauterine growth restriction.^3–6^

Polarity is a particularly important characteristic of epithelial cells, like the ST. Apical-basal polarity is the maintenance of discrete structure and function of the apical and basal surface of a cell and is a hallmark feature of many epithelia. In particular, the apical surface is dominated by microvilli, which are membrane protrusions supported by a core bundle of actin filaments (F-actin).^7,8^ Critically, active maintenance of cell polarity is important to support microvillar structural integrity. For example, microvillar F-actin cross-linkage to the cell membrane by linker proteins from the ezrin, radixin and moesin (ERM) family must be maintained via phosphorylation.^7,9,10^ Microvilli functionally increase the surface area of a cell to facilitate gas and nutrient exchange, are a site of endocytosis and signal transduction, and may be the site of extracellular vesicle release in the ST.^11,12^ Importantly, disruption of ST microvillar structure has been reported in placentas from intrauterine growth restriction and preeclampsia pregnancies.^3,13–16^ However, no studies have directly examined the mechanisms regulating maintenance of ST polarity and microvilli.

Apical-basal polarity is governed by evolutionarily conserved protein complexes. One such functional module is the Par complex that includes the scaffolding proteins partitioning defective-3 (Par-3), partitioning defective −6 (Par-6), and a Ser/Thr kinase, atypical protein kinase C (aPKC). The Par complex localizes apically and maintains apical identity in multiple cell types.^17–19^ The Par complex ultimately localizes and activates aPKC isoforms. APKC-ι and aPKC-ζ are the two major aPKC isoforms found in humans. We have shown that three isoforms of aPKC are expressed in ST: aPKC-ι, aPKC-ζ, and a novel N-terminal truncated *PRKCZ* encoded isoform – aPKC-ζ III.^20^ APKC-ζ III lacks the N-terminal PB-1 domain required for Par-6 binding.^20^,^21^ Hence, it is unclear if it can be fully activated since interaction with binding partners via the PB-1 domain is thought to be required for the full activation of aPKC isoforms.^21^,^22^ Importantly, aPKC isoforms have both individual and redundant functions. APKC-ζ global knockout mice have no embryonic phenotype,^23^,^24^ whereas aPKC-λ/ι global knockout mice are embryonic lethal at ~E9 with severe growth restriction and placental maternal-fetal interface malformation.^25^,^26^ But, aPKC-ζ can partially compensate for aPKC-λ/ι knockout in mouse embryos,^23^ therefore consideration for isoform specific and total aPKC function is necessary. Bhattacharya *et.al* found that placental specific and global knockout of aPKC-λ/ι lead to a lack of labyrinthine zone development, the primary zone for maternal/fetal exchange in mice.^25^ They also found that knockdown of *PRKCI* in human trophoblast stem cells decreased their ability to fuse and form ST, revealing an important role for aPKC-ι in maternal-fetal interface formation.^25^ However, the function of all aPKC isoforms in the ST has not been addressed to date.

In this study we sought to identify whether aPKC isoforms regulate apical polarization and microvillar organization of the ST. Our study reveals that aPKC isoform activity and expression is critical for the maintenance of the ST apical surface structure, integrity, and function in a regionalized manner. Moreover, we show that TNF-α regulates ST aPKC-ι expression, and that TNF-α and loss of aPKC activity leads to regionalized loss of ST microvilli and apical membrane integrity due to the induction of non-canonical pyroptosis.

## Results

Our initial studies examining aPKC isoforms in the human placenta revealed an apparent lack of anti-aPKC-ι or anti-aPKC-ζ/ζ III signal accumulation at the ST apical membrane in early first trimester placental samples.^20^ In other cell types, apical localization is required for aPKC’s to regulate apical-basal polarity, as mentioned above. Thus, we first performed colocalization analyses using antibodies targeting aPKC-ι or aPKC-ζ isoforms (which we previously validated to recognize aPKC-ζ and aPKC-ζ III^20^) and an anti-ezrin antibody, a ST apical membrane marker in third trimester placenta. ^27^ As observed previously,^20^ the anti-aPKC-ι signal was weak and largely diffuse within the ST from 4-6 weeks gestation (Supplementary Fig. 1A). There was also a lack of consistent apical accumulation of anti-ezrin signal at the ST apical surface despite a highly accumulated and complex pattern of anti-β-actin signal in the apical region (Supplementary Fig. 1A). By 7-8 weeks gestation, both anti-aPKC-ι and anti-ezrin signal had strong regional apical accumulation with a significantly increased colocalization compared to 4-6 weeks gestation (Fig. 1B, Supplementary Fig. S1A). Similar apical localization and aPKC-ι:ezrin colocalization coefficients were quantified in 9-12 week and 37-40 week ST (Fig. 1 A,B). Apical accumulation of anti-aPKC-ζ signal followed a similar trend, with a significant increase in the apical aPKC-ζ:ezrin colocalization coefficient by 7-8 weeks gestation (Fig. 1D, Supplementary Fig. S1B) that did not vary significantly at 9-12 weeks or 37-40 weeks gestation (Fig. 1C-D).

**Figure 1:**
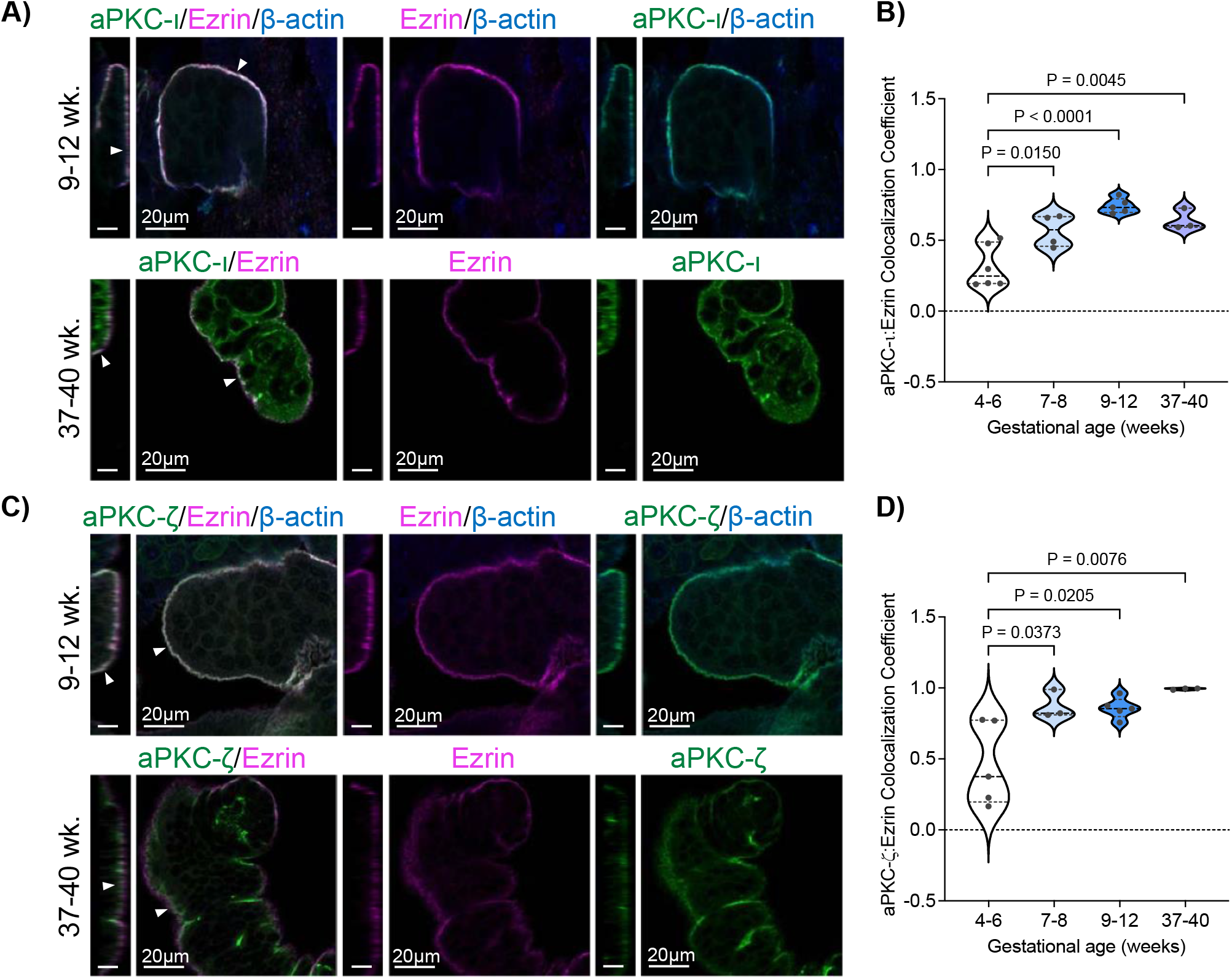
aPKC-ι and aPKC-ζ strongly colocalize with ezrin at the ST apical surface in 9-12 weeks and 37-40 week placenta. A) Representative images of 9-12 week (top panel) and 37-40 week (bottom panel) placenta tissue stained with anti-aPKC-ι (green), anti-ezrin (magenta) and anti β-actin [blue (top panels)] or anti-aPKC-ι (green) and anti-ezrin [magenta (bottom panels)]; left panel=zy plane, scale bar=10μm; right panel=xy plane; arrow heads=ST apical surface; single image plane of z-stack images; B) Summary data for quantitation of aPKC-ι and ezrin Pearson’s colocalization coefficient at the apical ST through varying gestational ages; *n*=3-6; bold dashed line=median; C) Representative images of 9-12 week (top panel) and 37-40 week (bottom panel) placental explants stained with anti-aPKC-ζ (green), anti-ezrin (magenta) and anti β-actin [blue (top panels)] or anti-aPKC-ζ (green) and anti-ezrin [magenta (bottom panels)]; left panel=zy plane, scale bar=10μm; right panel=xy plane; arrow heads=ST apical surface; single image plane of z-stack images; D) Summary data for quantitation of ST apical aPKC-ζ and ezrin Pearson’s colocalization coefficient through varying gestational ages; *n*=3-5; bold dashed line=median; All analyses performed with one-way ANOVA with Tukey’s post-hoc test.

Having established that aPKC isoforms and ezrin strongly and consistently colocalize at the ST apical surface from 9 weeks gestation, we used floating placental explant culture and a myristoylated aPKC pseudosubstrate inhibitor (aPKC inhibitor), which blocks both aPKC-ι and aPKC-ζ activity,^28^ to test if aPKC’s may regulate ST apical membrane structure via ezrin phosphorylation, as observed during murine intestinal development.^29^ After 6 hours of treatment, a significant decrease in the ST anti-phospho Thr-567 ezrin to total ezrin signal and the total anti-ezrin signal was observed in both 9-12 week and 37-40 week aPKC inhibitor vs. control treated explants (Fig. 2A-E, Supplementary Fig. S2). AKT was previously shown to regulate ezrin phosphorylation at Thr-567 in a trophoblastic cell line,^30^ so we also confirmed that aPKC inhibitor treatment did not alter the activity of AKT in 9-12 week placental explants (Supplementary Fig. S3A,B). Thus, aPKC activity regulates the ezrin Thr-567 phosphorylation and expression in ST. We also noted an appreciable decrease or change in signal pattern of apical anti-β-actin signal with aPKC inhibitor treatment (Fig. 2A). Therefore, we quantified the amount of ST apical F-actin in aPKC inhibitor treated explants using phalloidin. APKC inhibitor led to a greater than 50% decrease of ST apical phalloidin signal intensity in both first trimester and term explants (Fig. 2F-H, Supplementary Fig. S4A). Areas without appreciable changes in apical-phalloidin signal were interspersed with regions with clearly decreased signal. Significantly reduced apical phalloidin intensity was also observed after both 2 and 4 hours aPKC inhibitor treatment without the appearance of seemingly phalloidin-devoid regions (Supplementary Fig. S4B-D).

**Figure 2:**
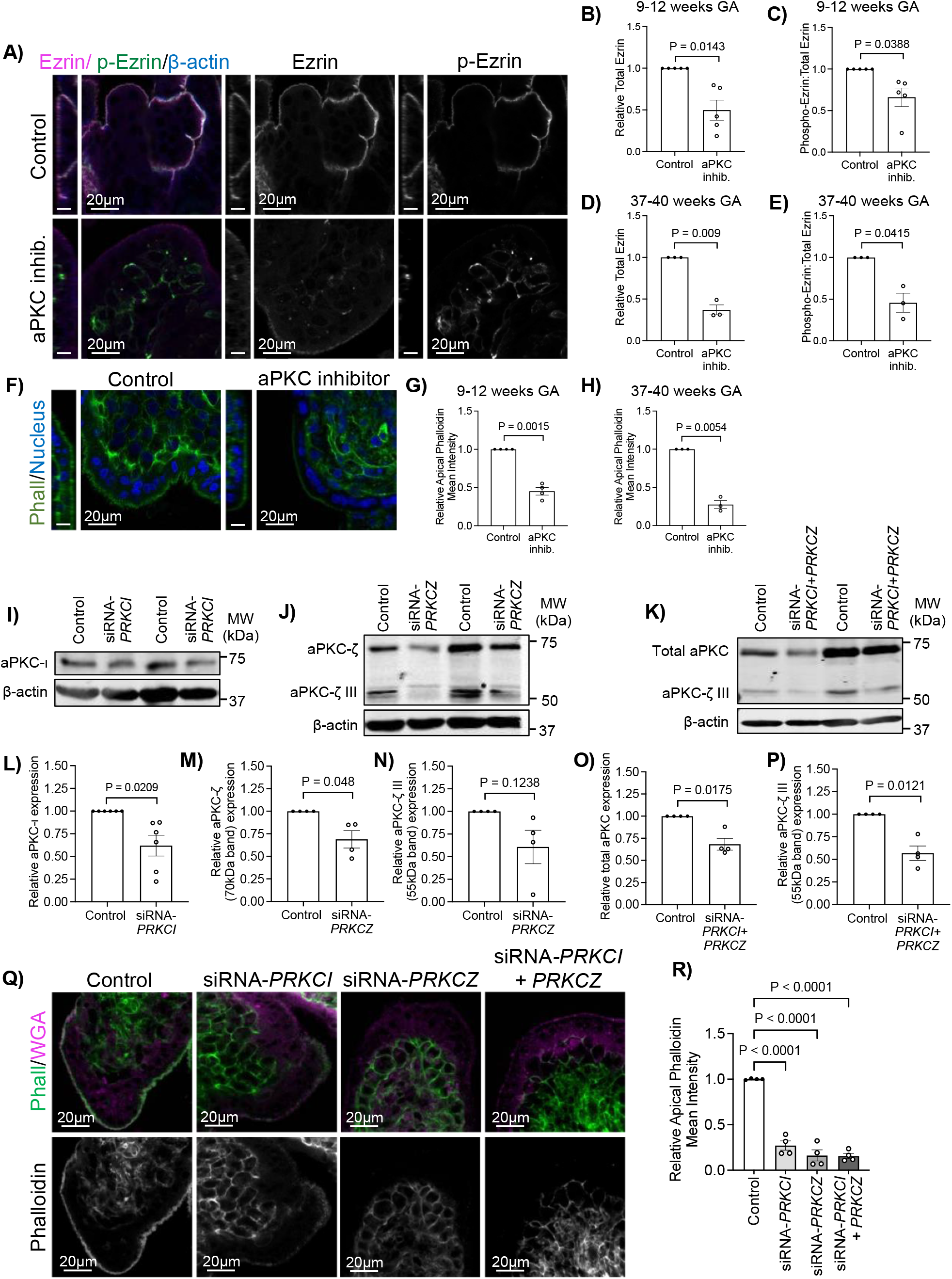
Loss of aPKC kinase activity or expression alters apical ezrin activation and abundance and F-actin. A) Representative images of 9-12 week placental explants treated +/- aPKC inhibitor for 6hrs stained with anti-phospho(Thr567)-ezrin (green), anti-ezrin (magenta) and anti-β-actin (blue); left panel=zy plane, scale bar=10μm; right panel=xy plane; single image plane of z-stack images; Summary data for quantitation of B) ST apical ezrin signal intensity in 9-12 week explants; *n*=5; C) ST apical phospho(Thr567) ezrin relative to total ezrin signal intensity in 9-12 week explants; *n*=5; D) ST apical ezrin signal intensity in 37-40 week explants; *n*=3; E) ST apical phospho(Thr567) ezrin relative to total ezrin signal intensity in 37-40 week explants; *n*=3; F) Representative images of phalloidin (green) and Hoechst-33342 (blue) in 9-12 week placental explants treated +/- aPKC inhibitor for 6hrs; left panel=zy plane, scale bar=10μm; right panel=xy plane; single image plane of z-stack images; Summary data for quantitation of ST apical phalloidin signal intensity in G) 9-12 week explants; *n*=4; H) 37-40 week explants; *n*=3; All above analyses performed using one-sample t-test; summary graphs mean +/- S.E.M; Representative western blot with I) anti-aPKC-ι and anti-β-actin following siRNA knockdown targeting *PRKCI;* J) anti-aPKC-ζ and anti-β-actin following siRNA knockdown targeting *PRKCZ;* K) anti-total-aPKC and anti-β-actin following siRNA knockdown targeting *PRKCI* and *PRKCZ;* Summary data for quantitation of relative L) aPKC-ι expression following siRNA knockdown targeting *PRKCI; n=6;* M) aPKC-ζ expression following siRNA knockdown targeting *PRKCZ; n*=4; N) aPKC-ζ III expression following siRNA knockdown targeting *PRKCZ; n*=4; O) total aPKC expression following siRNA knockdown targeting *PRKCI* and *PRKCZ; n=4;* P) aPKC-ζ III expression following siRNA knockdown targeting *PRKCI* and *PRKCZ; n*=4; L-P statistics= one-sample t-test; Q) Representative images of phalloidin (green) and WGA lectin (magenta) in 9-12 week placental explants treated with isoform specific siRNA for 24 hrs; left panel=zy plane, scale bar=10μm; right panel=xy plane; single image plane of z-stack images; R) Summary data for quantitation of ST apical phalloidin signal intensity in 9-12 week explants treated with aPKC isoform specific siRNA; *n*=3; Analysis performed using one-way ANOVA with Dunnett’s post-hoc test; summary graphs mean +/- S.E.M.

To address aPKC isoform-specific roles in ST, we performed siRNA mediated knockdown. 9-12 weeks explants were treated with siRNA targeting *PRKCI* and/or *PRKCZ*. Knockdown efficiency for aPKC-ι, aPKC-ζ, and aPKC-ζ III was determined by western blot analyses of explant lysate (Fig. 2I-P). Treatment with *PRKCI* siRNA, *PRKCZ* siRNA, or both lead to significantly decreased ST apical phalloidin mean intensity (Fig. 2Q-R). Importantly, as with aPKC inhibitor treatment, the appearance of seemingly-phalloidin replete areas was regionally specific with siRNA knockdown. Additionally, no change in the thickness of the ST could be observed using wheat germ agglutinin (WGA-lectin; Fig. 2Q), a lectin that binds to the apical surface and cytoplasm of the ST.^31,32^ A significant decrease in apical F-actin was observed with individual application of *PRKCI* siRNA and *PRKCZ* siRNA and no additive effect was observed with the addition of both. Similar results were achieved with a second set of siRNAs targeting *PRKCI* or *PRKCZ* (Supplementary Fig. 5). This suggests that reduced expression of a single isoform is sufficient to alter the ST apical actin cytoskeletal dynamics and that aPKC isoforms redundantly regulate ST apical membrane F-actin abundance.

To confirm if loss of aPKC kinase activity leads to alterations in the ST apical membrane structure we performed electron microscopy on control and aPKC inhibitor treated first trimester placental explants. Scanning electron micrographs (SEM) revealed a severe loss of microvilli at the ST apical surface with inhibitor treatment, apparent coalition of the few remaining microvilli and a porous appearance of the apical membrane (Fig. 3A, Supplementary Fig. 6) in a region-specific manner. Transmission electron microscopy micrographs (TEM) confirmed the simplification of the apical surface structure apparent by SEM and revealed a ST specific loss of cytoplasmic density, a high abundance of variably-sized membrane coated vesicles, and swollen mitochondria (Fig. 3B). Underlying cytotrophoblast progenitor cells did not have appreciable changes in any of these parameters and the basement membrane thickness between the cells did not vary between treatment groups.

**Figure 3:**
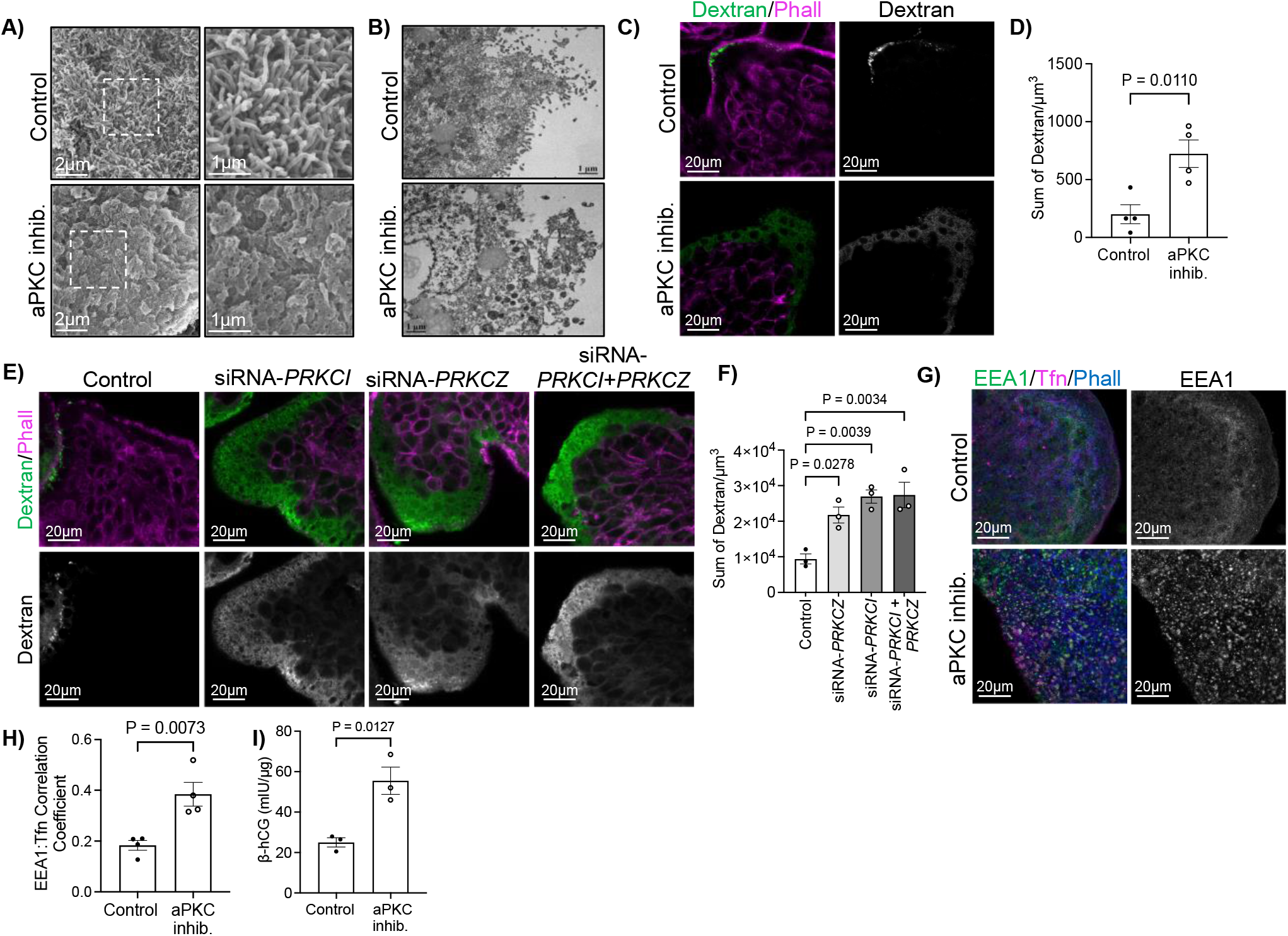
aPKC inhibition leads to ST apical membrane simplification, decreased microvilli and cytoplasmic density, permeabilization of ST, and alteration of ST endocytic trafficking. A) Representative SEM images of 9-12 week placental explants treated with aPKC inhibitor for 6hrs (representative of *n*=3); right panels=isolated zoomed images of boxed area indicated on left; B) Representative TEM images of 9-12 week placental explants treated +/- aPKC inhibitor for 6hrs (representative of *n*=3); C) Representative images of 9-12 week placental explants dextran-Texas Red (green) and phalloidin (magenta) following 2hrs treatment with aPKC inhibitor; left panels=merged image; right panels=isolated dextran signal; single plane of z-stack images; D) Summary data for quantitation of sum of dextran signal in 9-12 week placental explants after 2hrs aPKC inhibitor treatment; *n*=4; Student’s t-test; E) Representative images of 9-12 week placental explants dextran-Texas Red (green) and phalloidin (magenta) following 24hrs treatment with scramble control or siRNA sequences targeting *PRKCI* and/or *PRKCZ;* top panels=merged image; bottom panels=isolated dextran signal; single image plane of z-stack images; F) Summary data for quantitation of sum of dextran signal in 9-12 week placental explants following siRNA knockdown of *PRKCI* and/or *PRKCZ*; *n*=3; one-way ANOVA with Dunnett’s post-hoc test; G) Representative images of anti-EEA1 (green), transferrin-594 (magenta), and phalloidin (blue) in 9-12 week placental explants after 2hrs aPKC inhibitor treatment; top panels=control; bottom panels=aPKC inhibitor; right panels=isolated EEA1 signal; single plane of z-stack image H) Summary data for quantitation of Global Pearsons correlation coefficient for EEA1:transferrin in 9-12 week placental explants; *n*=4; Student’s t-test; I) Summary data for quantitation of β-hCG concentration normalized to total protein levels in explant conditioned medium +/- aPKC inhibitor treatment; *n*=3, Student’s t-test; All summary graphs mean +/- S.E.M.

Since SEM and TEM imaging revealed a regionalized decrease in cytoplasmic density and an almost lace-like appearance in the apical membrane we hypothesized that there is an increased permeability in ST with inhibition of aPKC’s. Indeed, aPKC inhibitor treatment and *PRKCI+PRKCZ* siRNA knockdown both lead to a ~3.5-fold increase in the uptake of 10,000 MW neutral dextran (Fig. 3C-F) compared to controls. *PRKCI* and *PRKCZ* siRNA treated samples also displayed a significant *3.1* and 2.5-fold increase in sum dextran signal (Fig. 3E,F). Importantly, this strong diffuse pattern of dextran uptake was restricted to areas of the ST lacking a visible continuous apical phalloidin signal. Thus, decreased aPKC isoform kinase activity or expression disrupts ST apical membrane integrity and permeabilizes the ST to a neutrally charged 10,000 MW compound.

In addition to the effects on the apical membrane, TEM also revealed the appearance of highly variably sized cytoplasmic membrane coated vesicles (Fig. 3B). In combination with the data showing altered F-actin abundance we hypothesized that loss of aPKC activity may lead to disrupted ST endocytic trafficking. Interestingly, we saw an increase in the colocalization of fluorescently-conjugated transferrin with anti-early-endosome antigen-1 (EEA1) when a transferrin uptake assay was performed on first trimester explants, suggesting an increase in clathrin-mediated transferrin endocytosis with aPKC inhibitor treatment (Fig. 3G,H; Supplementary Fig. 7A-E). There were also enlarged areas of anti-EEA1 signal at the ST apical surface in first trimester explants treated with aPKC inhibitor (Fig. 3G; Supplementary Fig. 7B,D). Interestingly, these enlarged EEA1 signals were not exclusively associated with regions of the ST with visibly decreased phalloidin signal. These data suggest that inhibition of aPKC’s dysregulates ST clathrin-mediated endocytosis and leads to an increased endocytic activity or stalling of endosomal trafficking in the apical compartment.

We reasoned that the altered vesicular trafficking and increased cellular permeability induced by aPKC inhibitor treatment may also lead to increased release of ST-derived factors. The ST produces several pregnancy-sustaining hormones, including the hormone human chorionic gonadotropin (hCG). Therefore, we quantified the release of the β-hCG subunit via ELISA in explant conditioned medium from control and aPKC inhibitor treated samples. As expected, aPKC inhibitor led to a nearly 3-fold increase of β-hCG in explant conditioned medium (Fig. 3I). In summary, aPKC isoforms regulate the integrity of the apical membrane, vesicular trafficking within and structure of the apical compartment, likely via the regulation of F-actin and ezrin abundance.

TNF-α has been shown to decrease expression of aPKC-ι,^33^ and is known to be elevated in the maternal circulation in established placental pathologies^34,35^ where regionalized loss of microvilli, reduced cytoplasmic density, and hyper-vesicular cytoplasm have also been observed in the ST.^3,14,16^ Hence, we hypothesized that TNF-α would also decrease the expression of aPKC isoforms in the ST. Interestingly, we found that treatment of *in vitro* differentiated primary ST with TNF-α led to a dose dependent significant decrease in aPKC-ι expression only (Fig. 4A-D). Additionally, both 100pg/mL and 10ng/mL doses of TNF-α resulted in a regionalized but overall significant decrease in apical phalloidin in both first trimester and term explants (Fig. 4E-G, Supplementary Fig. 8A), like aPKC-inhibitor treatment or aPKC isoform siRNA knockdown. As expected, SEM revealed a regionalized severe loss of microvilli at the ST apical surface with TNF-α treatment and a similar porous or lace-like appearance of the apical membrane (Fig. 4H; Supplementary Fig. S8A) like with aPKC-inhibitor treatment. To confirm if this led to ST permeabilization, we performed dextran uptake assays that revealed a diffuse and significantly increased dextran signal in ST compared to control in both first trimester and term TNF-α-treated explants (Fig. 4I-K, Supplementary Fig. 8B). Additionally, when explants were imaged by brightfield microscopy for 24 hours discrete areas of membrane blebbing on the placental surface were seen starting 12 hours after treatment with TNF-α (Fig. 4L; Supplementary Movies 1,2). Thus, TNF-α leads to isoform specific dose dependent decrease in ST aPKC-ι expression as well as simplification, permeabilization, and blebbing of the ST apical surface.

**Figure 4:**
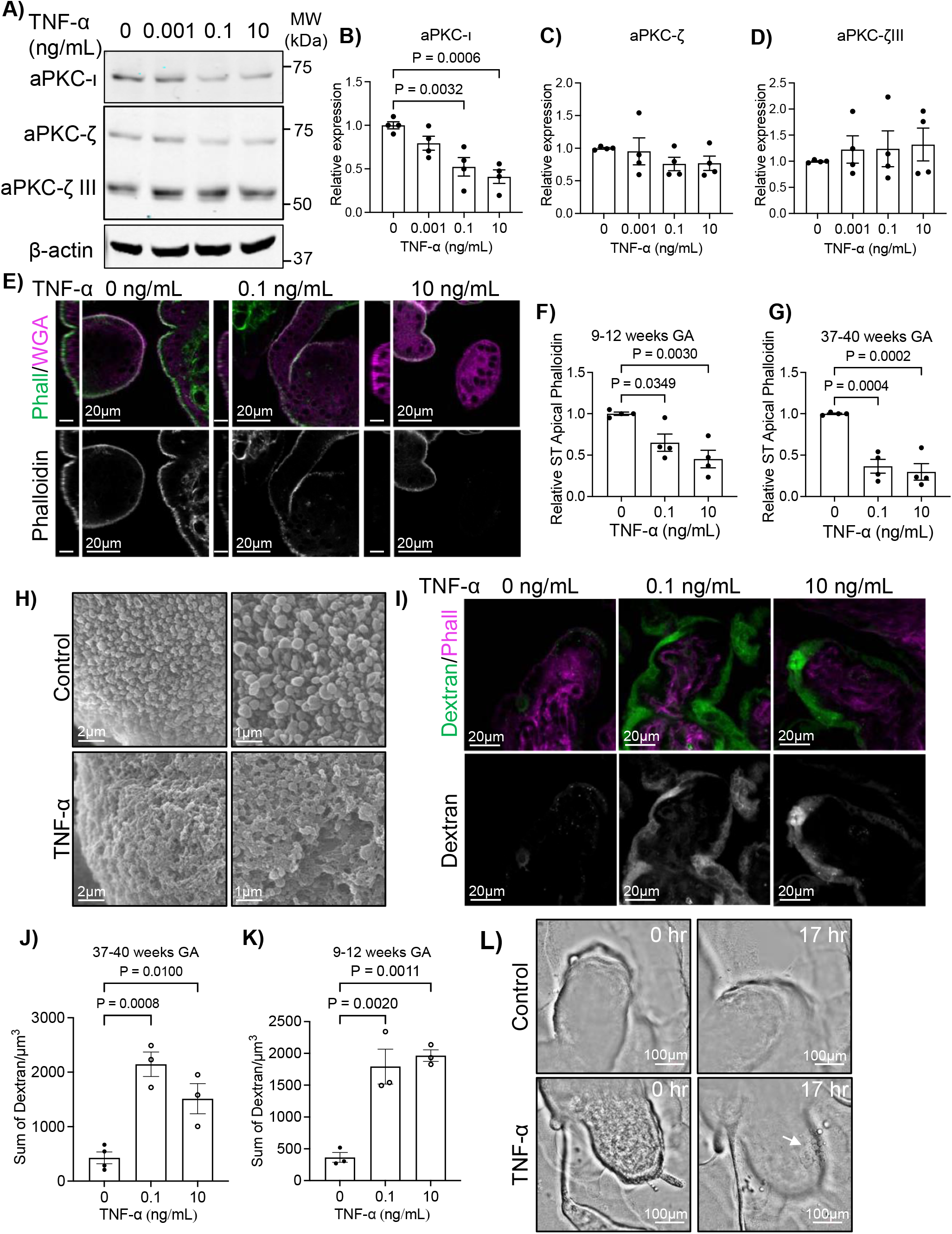
TNF-α leads to dose dependent isoform specific decrease in aPKC-ι expression, loss of apical F-actin, and permeabilization of ST. A) Representative western blot analyses of *in vitro* differentiated primary ST treated for 4hrs with indicated doses of TNF-α, immunoblotted with anti-aPKC-ι, anti-aPKC-ζ, and anti-β-actin; B) Western blot quantitation for aPKC-ι; C) aPKC-ζ (70kDa form); and D) aPKC-ζ III (55kDa form); *n*=4 patient derived cells; E) Representative images of phalloidin (green) and WGA lectin (magenta) in 9-12 week placental explants treated with indicated doses of TNF-α for 6hrs; top panels=merged image; bottom panels= isolated phalloidin; left panel=zy plane, scale bar=10μm; right panel=xy plane; single image plane of z-stack images; F) Summary data for quantitation of ST apical phalloidin signal intensity; *n*=4; G) Summary data for quantitation of ST apical phalloidin signal intensity; *n*=4; H) Representative SEM images of 37-40 week explants +/- 100 pg/mL TNF-α for 6hrs; right panels= higher magnification images of the same samples; *n*=4; (I) Representative images of dextran-Texas Red (green) uptake and phalloidin (magenta) in 37-40 week placental explants after 6hrs treatment with indicated doses of TNF-α; top panels=merged image; bottom panels=isolated dextran; single plane image of z-stack (J) Summary data for quantitation of sum dextran in the ST; *n*=3; (K) Summary data for quantitation of sum dextran signal in the ST; *n*=3; (L) Representative images of 9 week placental explants at 0 hours and 17 hours treatment +/- 100pg/mL TNF-α; arrow indicate regions of membrane blebbing; All analyses one-way ANOVA with Dunnett’s post-hoc test; summary graphs mean +/- S.E.M.

Loss of aPKC expression/activity or exposure to TNF-α induce multiple forms of cell death, including apoptosis and pyroptosis, in other cell types.^36–42^ Pyroptosis is a pro-inflammatory form of cell death characterized by the appearance of membrane pores due to the cleavage and oligomerization of gasdermin family proteins.^40^,^43^ Gasdermin pore formation leads to the permeabilization of the membrane to low molecular weight compounds,^44^ and the release of mature IL-1β and other damage associated molecular patterns (DAMPS).^43^,^45^,^46^ Since TNF-α and aPKC inhibitor treatment both induced regionalized formation of pore-like structures and increased permeability we hypothesized that they were inducing ST pyroptosis. Interestingly, when IL-1β levels were tested in explant conditioned medium a significant increase was only observed in first trimester explant-conditioned medium, but not term explant conditioned medium for both aPKC inhibitor and TNF-α (Fig. 5A-D). The ST has previously been shown to express gasdermin D,^47^ therefore we stained TNF-α treated explants with an anti-cleaved (Asp275) gasdermin D (cl-GSDMD) antibody expecting to see an increase in cl-GSDMD signal at the ST apical membrane. In contrast to our hypothesis, no anti-cl-GSDMD signal could be observed within the ST, despite clear signal in both control and TNF-α treated cells in the villous core (Fig. 5E). A similar staining pattern was observed in aPKC inhibitor treated explants (Supplementary Fig. 9A). No cl-GSDMD signal was detected in tissue that was fixed without culture whereas a stromal cl-GSDMD signal was observed in explant cultured tissue from the same donor (Supplementary Fig. 10). Gasdermin E is a second gasdermin family member known to be expressed in the placenta.^43^ We first confirmed that it is expressed in the ST at both 9-12 weeks and 37-40 weeks gestation using an anti-gasdermin E antibody (Fig. 5F-G; Supplementary Fig. 11A-C), where a punctate signal pattern was observed in discrete areas of the ST, as well as sporadic cytotrophoblast progenitor cells at all gestational ages examined (Supplementary Fig. 10B). Interestingly, TNF-α treatment led to an apparent aggregation of anti-GSDME signal into large high signal intensity puncti at the apical border of the accumulated dextran signal in first trimester explants (Fig. 5F). A similar change in anti-gasdermin E signal was also observed in aPKC-inhibitor treated first trimester explants (Supplementary Fig. 9B). In term explants only rare large aggregates of anti-gasdermin E signal observed in the apical region, though high-signal intensity aggregates were observed in the basal region of the ST with both treatments (Fig. 5G, Supplementary Fig. 8C). To confirm if gasdermin E cleavage occurs in the ST *in vitro* differentiated ST were treated with TNF-α and prepared for western blotting analysis. Both full-length gasdermin E and p30 cleaved gasdermin E bands were observed, with a 2-fold increase in p30 gasdermin E in TNF-α treated cells (Fig. 5H-I). Caspase-3 has been shown to cleave gasdermin E, allowing for oligomerization and pore formation,^42,48^ therefore we used an anti-active caspase-3 antibody (cleaved caspase-3) to stain TNF-α and aPKC inhibitor treated explants. No signal could be detected within the ST in any treatment group in both first trimester and term explants (Fig. 5J,K; Supplemental Fig. 9D,E), though a clear signal could be detected in stromal cells, especially in TNF-α treated explants, consistent with previous studies showing caspase-3 activity is restricted to the cytotrophoblasts.^49^

**Figure 5:**
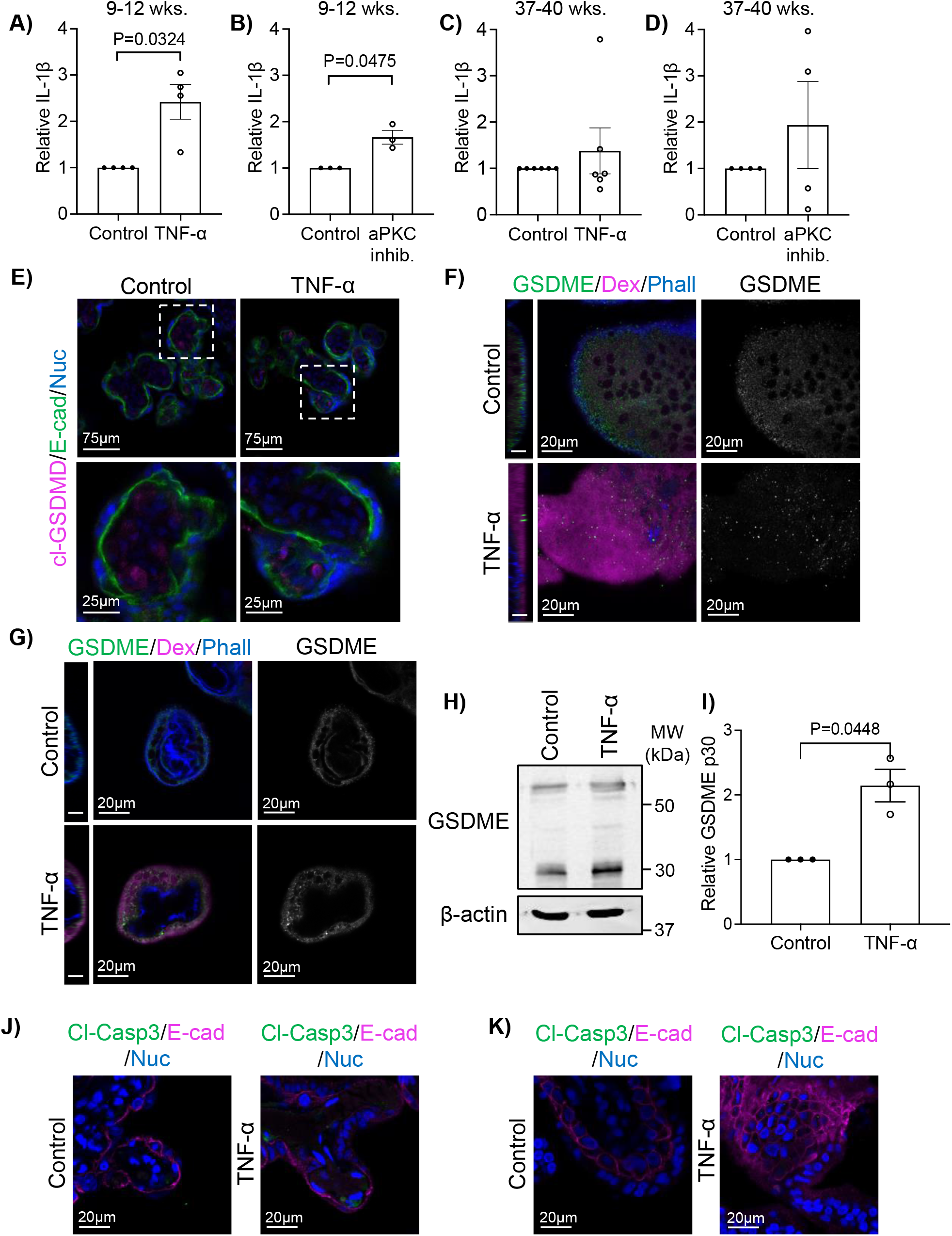
TNF-α leads to cleavage of gasdermin E. A) IL-1β in placental explant medium after 6hr 100pg/mL TNF-α stimulation; *n*=4; B) IL-1β placental explant medium after 6hr aPKC inhibitor treatment; *n*=3; C) IL-1β in placental explant medium after 6hr 100pg/mL TNF-α stimulation; *n*=6; D) IL-1β in placental explant medium after 6hr aPKC inhibitor treatment; *n*=4; All analyses (A-D) one-factor t-test; E) Representative image of anti-cleaved gasdermin D (cl-GSDMD, magenta); anti-E-cadherin (green), and Hoescht 33342 stained 37-40 week placental explants +/- 100pg/mL TNF-α for 6hrs; bottom panels= higher magnification images of indicated areas; single xy image plane; F) Representative images of anti-gasdermin E (GSDME, green), dextran-Texas Red (magenta), and phalloidin (blue) in 9-12 week placental explants treated +/- 100pg/mL TNF-α for 6hrs; left panel=yz single image plane of merged image scale bar=10μm; center panel= xy single image plane of merged image; right panel=gasdermin E alone xy single image plane; G) Representative images of anti-gasdermin E (GSDME, green), dextran-Texas Red uptake (magenta), and phalloidin (blue) in 37-40 week placental explants treated +/- 100pg/mL TNF-α for 6hrs; left panel=yz single image plane of merged image; scale bar=10μm; center panel=xy single image plane of merged image; right panel= gasdermin E alone; xy single image plane; H) Representative western blot of *in vitro* differentiated primary ST treated +/- 10ng/mL TNF-α for 12 hrs; I) Quantitation of western blot analyses for gasdermin E p30 normalized to total protein; *n*=3 experimental replicates from *n*=2 patient derived cells; one-factor t-test; J) Representative images of anti-cleaved caspase 3 (cl-Casp3, green), anti-E-cadherin (magenta), and Hoescht 33342 stained 37-40 week placental explants +/- 100pg/mL TNF-α for 6hrs; K) Representative images of anti-cleaved caspase 3 (cl-Casp3, green), anti-E-cadherin (magenta), and Hoescht 33342 stained 9-12 week placental explants +/- 100pg/mL TNF-α for 6hrs; All graphs mean +/- S.E.M.

To identify if gasdermin E is necessary for the increased permeability to dextran observed with TNF-α and aPKC inhibitor treatment, 9-12 week explants were treated with gasdermin E targeting siRNA and then stimulated with either TNF-α and aPKC inhibitor. A significant decrease in gasdermin E expression was observed after 48hrs treatment with siRNA by western blotting analysis (Supplemental Fig. 12). Critically, gasdermin E knockdown blocked increased dextran permeability induced by both TNF-α and aPKC inhibitor (Fig. 6A-D). Dimethyl fumarate (DMF) is an anti-pyroptotic agent that blocks the cleavage of both gasdermin D and -E via succination.^50^ Therefore, we also sought to block TNF-α and aPKC inhibitor-induced pyroptosis by co-administration of the factors with DMF. As expected, DMF prevented TNF-α induced ST permeability to dextran in both first trimester and term explants (Fig. 6E,F; Supplemental Fig. 13). First trimester aPKC-inhibitor induced dextran accumulation could also be blocked by DMF (Fig. 6 G,H). Hence, the sum of these data reveal that TNF-α and loss of aPKC kinase activity leads to the induction of ST pyroptosis, via a gasdermin E mediated pathway.

**Figure 6:**
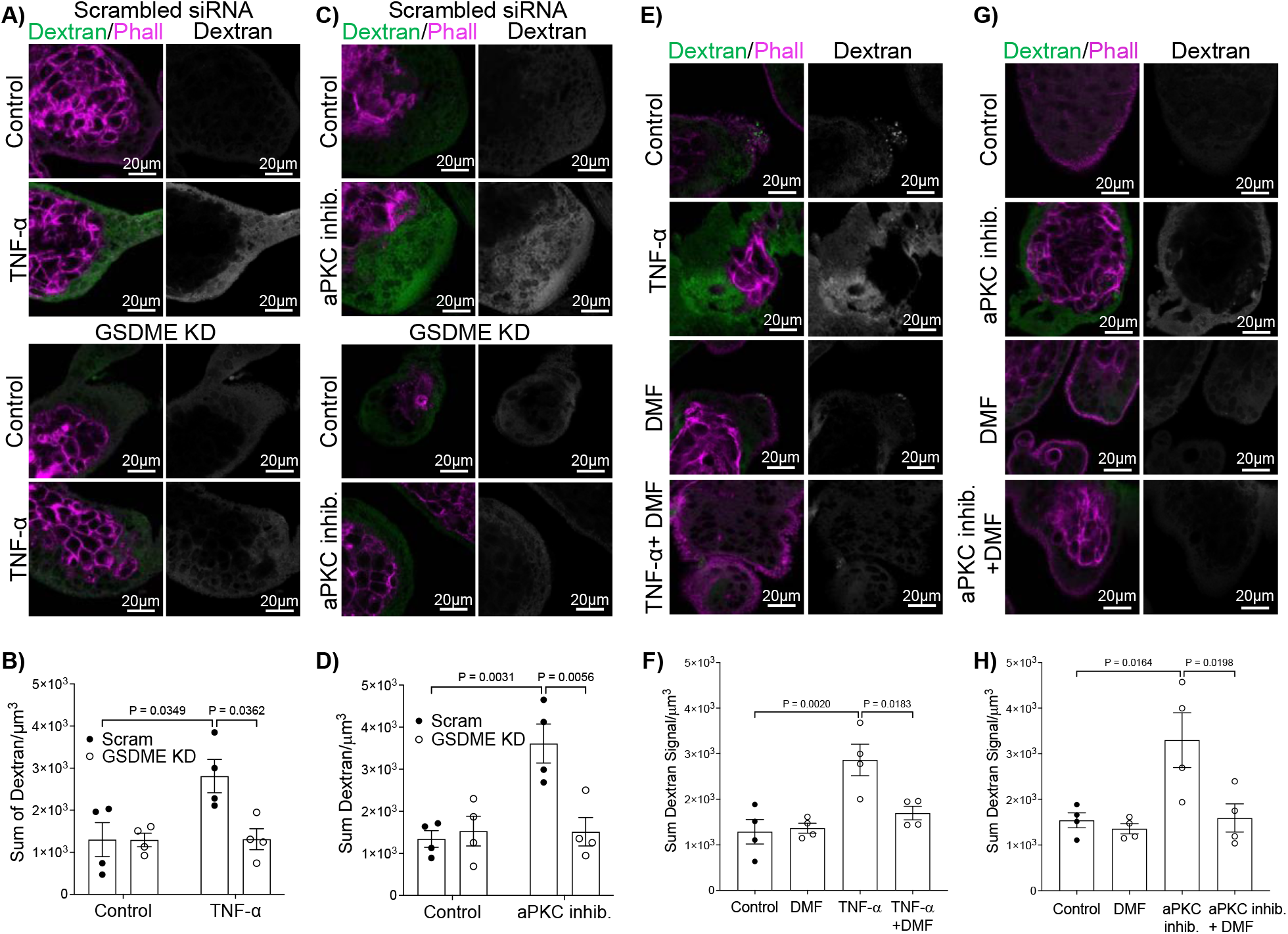
GSDME knockdown blocks TNF-α and aPKC inhibitor induced ST permeability. A) Representative images of dextran-Texas Red (green) and phalloidin (magenta) in 9-12 week placental explants +/- GSDME siRNA and 6hrs treatment +/- 100pg/mL TNF-α; B) Summary data for quantitation of sum dextran signal in the ST of 9-12 week placental explants; *n=4;* C) Representative images of dextran-Texas Red (green) and phalloidin (magenta) in 9-12 week placental explants +/- GSDME siRNA and 6hrs treatment +/- aPKC inhibitor; D) Summary data for quantitation of sum dextran signal in the ST of 9-12 week placental explants; *n*=4; E) Representative images of dextran-Texas Red (green) and phalloidin (magenta) in 9-12 week placental explants after 6hrs treatment +/- DMF, 100pg/mL TNF-α, or both; F) Summary data for quantitation of sum dextran signal in the ST of 9-12 week placental explants; *n*=4; G) Representative images of dextran-Texas Red (green) and phalloidin (magenta) in 9-12 week placental explants after 6hrs treatment +/- DMF, aPKC inhibitor, or both; H) Summary data for quantitation of sum dextran signal in the ST of 9-12 week placental explants; *n*=4; B= two-way ANOVA with Sidak’s multiple comparison test; D= two-way ANOVA with Tukey’s multiple comparison test; F,H= one-way ANOVA with Sidak’s multiple comparison test; All graphs mean +/- S.E.M.; All images are a single xy image plane.

## Discussion

The ST inhabits an exceptional anatomical position as a fetal-derived cell bathed in maternal blood at its apical surface but attached to the fetal compartment at its basal edge. Here we found that aPKC isoforms regulate the structure, permeability, and function of the ST apical surface in a spatially restricted manner by controlling the initiation of gasdermin E dependent pyroptosis. Additionally, we established that the pro-inflammatory cytokine TNF-α decreases ST aPKC-ι expression and profoundly alters ST apical structure and permeability, like disruption of aPKC isoforms, also leading to the initiation of pyroptosis. Therefore, our data suggests that aPKC isoforms are key regulators of ST homeostasis that can rapidly change in expression upon exposure to a pro-inflammatory stimulus leading to the release of a potent pro-inflammatory cytokine into maternal circulation.

Given the strong effects observed with the disruption of aPKC isoform expression and activity and that TNF-α leads to the isoform specific decrease in aPKC-ι expression it will be important to identify the relevant regulators controlling aPKCs within the ST. It is presently unknown whether aPKC isoforms function as a part of the Par-polarity complex in this cell type. Par-6 expression at the ST apical membrane from 12-weeks gestation has been shown,^51^ which coincides with the gestational age range we observed a significant signal accumulation for all aPKC isoforms at the apical surface (Fig. 1), but the expression and localization of Par-3 within the ST is unknown. Our data revealed profound regional disruption of the apical F-actin cytoskeleton, but also the appearance of very large EEA1 positive vesicular structures in regions where F-actin was not visibly diminished. Therefore, there are likely multiple pathways through which aPKCs facilitate ST function to be identified. Similarly, aPKC inhibitor treatment significantly decreased the ratio of activated to total ezrin as well as total ezrin signal (Fig. 2), indicating that aPKC kinase activity could be directly phosphorylating ezrin at the Thr-567 residue as previously demonstrated,^29^ as well as indirectly controlling ezrin abundance via an additional pathway, to regulate ezrin homeostasis, microvillar structure, and maintenance. Intestinal cell regulation of aPKC-ι expression downstream of TNF-α is controlled by post-transcriptional mechanisms and ubiquitin-mediated degradation. Understanding how this is achieved in ST and whether other pro-inflammatory cytokines have similar effects will also be important in the future.

Our work also demonstrates that ST undergoes pyroptosis via a gasdermin E mediated pathway. These results are consistent with multiple studies that have identified ST features by electron microscopy interpreted as regionalized necrosis.^3,16^ These microvilli-replete ST regions have especially been identified in ST from preeclamptic pregnancies,^16^ where increased placental mature IL-β has also been reported.^52^ Placental mature-IL-1β was found to be predominantly produced by the ST^47^ where the authors also observed a lack of cleaved-gasdermin D signal, thereby supporting our conclusions that ST pyroptosis is not mediated by gasdermin D. Though, it is presently unclear how gasdermin E cleavage is occurring since no detectable caspase-3 activation was seen in ST in our experiments. The lack of caspase-3 activation is consistent with previous literature showing that cleaved-caspase-3 is restricted to the cytotrophoblast progenitors *in vivo.^49^* Additionally, it is presently unknown how aPKC inhibition stimulates gasdermin E-dependent pyroptosis in the ST. Therefore, further elucidation of the ST pyroptotic cascade is necessary.

Our data showing that the release of IL-1β was not significantly altered by TNF-α treatment or aPKC inhibitor in term placental explants indicates that gestational-age dependent mechanisms blunting the release of pro-inflammatory factors from the ST may exist, despite a significant permeabilization of the ST to dextran. This is consistent with the observations of Megli *et al*. who found that constitutive NLRP3 inflammasome activation in the ST was significantly dampened at term compared to second trimester explants.^47^ Elucidating additional pyroptotic-initiating factors, and whether the mechanism of action has conserved regulatory points that could serve as therapeutic targets to block ST pyroptosis will also be an interesting future direction.

Finally, our work has clear implications for the pathobiology of placental disorders and infection during pregnancy. Presently there is no data that we are aware of examining aPKC isoform expression in the placentas from pregnancy complications. Our results revealed that disruption of aPKC isoforms induces the rapid appearance of features characteristic of regions of the ST from preeclamptic pregnancies. Though preeclampsia is defined by the onset of maternal hypertension after the 20^th^ week of gestation and end organ failure,^53–55^ it is appreciated that the pathologic processes necessary for the development of the most severe form of the syndrome, early-onset preeclampsia, occur in early gestation.^4^ Interestingly, increased maternal first trimester circulating levels of IL-1β have been reported in pregnancies that go on to develop early-onset preeclampsia.^56^ Therefore, our data showing that pyroptosis can be initiated in 9-12 week placenta suggest that chronic initiation of this pathway could contribute to the progression of some forms of preeclampsia and should be directly tested in the future. We also showed that regionalized membrane blebbing occurs during TNF-α stimulated ST pyroptosis in first trimester placental explants. An increase in ST derived microvesicles in maternal circulation of women with preeclampsia has been shown and is postulated to be a major mechanism leading to the development maternal vascular dysfunction in the syndrome.^57,58^ Our work suggests that ST pyroptosis may be the cause of increasing circulating levels of microvesicles, which could begin as early as the end of the first trimester. Therefore, thorough characterization of the size, quantity, and type of vesicles released from the pyroptotic ST is another important future direction.

## Materials and Methods

### Tissue collection

37-40 weeks gestation and first trimester human placental samples were collected by methods approved by the University of Alberta human research ethics board (Pro00088052, Pro00089293). Term placental tissue was collected after cesarean delivery without labour from uncomplicated pregnancies. First trimester placental tissue was obtained from elective pregnancy terminations following informed consent from the patients.

### Floating placental explant culture and treatments

Placental samples were collected and rinsed in cold 1X phosphate-buffered saline (PBS) to remove blood. For term placentas, tissue was cut from three central cotyledons, decidua and blood clots were removed, and trimmed tissue was washed extensively in PBS to remove residual blood. For first trimester samples, placenta was identified, separated from decidua, blood clots were removed, and tissue was washed extensively in PBS to remove residual blood. Both first trimester and term tissue were then cut into uniform 2 mm^3^ explants, placed into 48-well plates and incubated overnight at 37°C 5% CO_2_ in Iscove’s Modified Dulbecco’s Medium (IMDM; Gibco; Grand Island NY) supplemented with 10% (v/v) Fetal Calf Serum (FCS; Multicell, Wisent Inc.; Saint-Jean Baptiste Canada) and penicillin streptomycin (5000 IU/mL; Multicell, Wisent Inc.). Following overnight incubation, explants were washed in serum-free IMDM with 0.1% (w/v) bovine serum albumin (BSA, Sigma-Aldrich; St. Louis MO) and incubated for 2-24hrs at 37°C 5% CO_2_ +/- myristoylated aPKC pseudosubstrate inhibitor (5 μM; Invitrogen; Product number 77749) in IMDM+0.1% BSA. For TNF-α treatments, after washing in serum free medium, explants were pre-incubated for 30 min at 37 °C 5%CO_2_ in IMDM +0.1% BSA, then medium was changed to IMDM +0.1% BSA +/- TNF-α (100 pg/mL or 10 ng/mL; Sigma-Aldrich) and incubated at 37°C 5% CO_2_ for 6-24hrs. Explants were then washed with cold PBS and fixed with 4% paraformaldehyde for 2 hrs on ice. For experiments performed with dimethyl fumarate (DMF; Sigma Aldrich), explants were washed as above, and then pre-incubated for 30 minutes with solvent control (DMSO; 1:1000) or DMF (25μM). Medium was then changed and 100pg/mL TNF-α or 5 μM myristoylated aPKC pseudosubstrate inhibitor was added +/- DMF and incubated for 6hrs at 37°C 5% CO_2_.

Explants were then washed with cold PBS and fixed with 4% paraformaldehyde for 2 hrs on ice. Technical triplicates were performed for all treatments.

### siRNA Knockdown

siRNA knockdown was performed as previously reported.^20^ 9-12 week placental explants were placed into a 48-well plate with IMDM supplemented with 10% (v/v) FCS and gentamicin (50 μg/mL; Gibco). siRNA sequences targeting *PRKCI* (final concentration 0.2 μM; ON-TARGETplus siRNA J-004656-07or J-04656-10; Dharmacon; Lafayette CO) and *PRCKZ* (final concentration 0.2 μM; ON-TARGETplus siRNA J-003526-17-0010 or J-003526-14; Dharmacon), siRNA targeting both *PRKCI* and *PRCKZ*, or scrambled control (final concentration 0.2 μM; ON-TARGETplus Non-targeting Control Pool D-001810-10-20; Dharmacon) were added to the medium and incubated for 24hrs. siRNA sequences targeting *DFNA5* (final concentration 0.2 μM; hs.Ri.DFNA5.13.1; Integrated DNA Technologies; Coralville, IA) or negative control (final concentration 0.2 μM; Integrated DNA Technologies) were added to the medium and explants were incubated for 48hrs. After treatment, explants were washed with cold PBS before fixation with 4% paraformaldehyde for 2hrs on ice or collected in RIPA lysis buffer to perform western blotting. For gasdermin E siRNA treated explants, aPKC inhibitor and TNF-α treatment was added for 6hrs 48hrs after culture with the siRNA.

### Dextran Uptake Assay

In the last 30 minutes of treatment with aPKC inhibitor, TNF-α, or siRNA explants were incubated with 10,000 molecular weight (MW) neutral Dextran Texas Red^™^ (25 μg/mL; D1828; Invitrogen, Eugene OR) for 25 minutes in IMDM+0.1% BSA and washed with cold PBS before fixation with 4% paraformaldehyde for 2hrs on ice.

### Transferrin endocytosis assay

Following aPKC inhibitor treatment, explants were incubated with fluorescently conjugated human transferrin-594 (CF 594; 25 μg/mL; Biotium Fremont CA) for 40 minutes. Explants were then washed extensively with cold 1X PBS and fixed in 4% paraformaldehyde for 2hrs on ice.

### Primary trophoblast isolation, culture, and treatment

Term placental cytotrophoblasts were isolated according to previously published methods.^20^,^59^ To obtain *in vitro* differentiated syncytiotrophoblasts, isolated cytotrophoblast progenitor cells were seeded into 6 well plates and cultured in IMDM+10%FCS+ Penicillin-streptomycin and incubated for 4hrs at 37°C 5%CO_2_. Cells were then washed to remove non-adherent cells, medium changed to IMDM+10%FCS+Penicillin-streptomycin + 8-bromo-cAMP (10μM, Sigma-Aldrich), and then incubated overnight at 37°C 5%CO_2_. Medium was changed to remove the 8-bromo-cAMP and the cells were incubated for a further 48hrs 37°C 5%CO_2_ (72hrs in culture total).

TNF-α treatments were performed after differentiation into syncytiotrophoblasts (after 72hrs in culture). Medium was removed, cells were washed, and medium was replaced with IMDM+0.1%BSA and cells were incubated for 30 min at 37°C 5%CO_2_. Medium was then changed to IMDM+0.1%BSA +/- TNF-α at the indicated doses for 4-12hrs. Cells were then washed, and protein lysates prepared for western blotting analysis.

### Western Blotting

Samples were prepared by adding RIPA lysis buffer and protease inhibitor (1:100; Sigma Aldrich; P2714) and protein concentration was determined using a BCA Protein Assay.

Protein was loaded and run on SDS-polyacrylamide gels before transfer onto nitrocellulose membranes. The membranes were probed with mouse anti-aPKC-ι (1:1000; BD Biosciences; 610207), rabbit anti-aPKC-ζ (1:2000; Atlas Antibodies; HPA021851), mouse anti-total aPKC (1:1000; Santa Cruz Biotechnologies; sc-17781), mouse anti-AKT(pan) (1:2000; Cell Signaling Technology #2920), rabbit anti-phospho Ser473 AKT (1:2000; Cell Signaling Technology #4060), rabbit-anti-gasdermin E (1:10000; Abcam ab215191) and mouse anti-β-actin (1:10000; Cell Signaling Technology #8457) and fluorescent secondary antibodies. Secondary antibodies included Alexa Fluor® donkey anti-mouse 680 (1:10,000; Invitrogen; A28183) and Alexa Fluor® donkey anti-rabbit 800 (1:10,000; Invitrogen; A21039). Total protein quantification was performed by staining membranes with Fast-Green FCF.^60^ All blots were scanned on a Licor Odyssey scanner and quantitation was performed using the Licor Imaging Software with target protein band intensity normalized to total protein.

### Tissue staining and image analysis

Following fixation cultured explant or fixed un-cultured placental tissue was whole mount stained. Tissue was incubated with blocking buffer [5% normal donkey serum and 0.3% Triton x100, 1% human IgG (Invitrogen) in PBS] followed by incubation with primary antibodies: anti-aPKC-ι (1:100; Atlas Antibodies; HPA026574); anti-aPKC-ζ (1:200, Atlas Antibodies, HPA021851); anti-ezrin (1:33; Invitrogen; PA5-18541); anti-phospho Thr567 ezrin (1:200; Invitrogen; PA5-37763); anti β-actin (1:250; Cell Signaling Technologies; #8457); anti-EEA1 (1:50; R&D Systems MAB8047); anti-cleaved GSDMD Asp275 (1:500; Cell Signaling Technologies #36425); anti-GSDME (1:50; Santa Cruz Biotechnology; sc393162); anti-cleaved caspase-3 Asp175 (1:400; Cell Signaling Technologies; #9661) anti-E-cadherin (1:400; R&D Systems; MAB18381) or biotinylated-WGA Lectin (1:1000; Vector Biolabs) overnight, then washed and incubated with the appropriate secondary antibodies (Alexa Fluor^™^, Invitrogen) and/or fluorescently conjugated phalloidin (1:400; iFluor 405 or iFluor594; AAT Bioquest, Sunnyvale CA). Hoechst 33342 (Invitrogen) was then added for 30 mins. Tissue was then washed and mounted with imaging spacers.

All images were captured with a Zeiss LSM 700 confocal microscope using a Zeiss 20x/0.8 M27 lens or a Zeiss 63x/1.4 Oil DIC M27 lens. 10-15 μm Z-stack images with a 1 μm step size were captured at 63x magnification. Three images per treatment were captured. Image capture was restricted to blunt-ended terminal projections with underlying cytotrophoblast progenitors in villi from first trimester and term samples.

### ELISA assays

Conditioned explant culture medium was collected from technical triplicates after 6hrs incubation with treatments and centrifuged at 12,000 rpm for 10 minutes and the supernatant aliquoted then stored at −20°C. Explant tissue was washed with PBS then flash frozen and stored at −80°C until total protein was extracted and determined by BCA assay, as above. ELISAs were performed using a β-hCG ELISA kit (EIA-1911; DRG International Springfield NJ) and IL-1β ELISA kit (DY401-05, R&D Systems Minneapolis MN). Plates were read using the Biotek Synergy HTX plate reader (Gen 5 software). The β-hCG and IL-1β values were interpolated using GraphPad PRISM 9 (Version 9.3.1) and normalized to the total protein from the explant the medium was produced by.

### Live cell imaging

9-12 week placental explants were washed and cut as detailed above. Tissue was then immersed in 1% low melting point agarose (Sigma) at 40°C for less than 30 seconds, placed in a 48 well plate and overlaid with 5μL of additional 1% low melting point agarose. Agarose was allowed to cool for 15 min at room temperature, then IMDM+10% FCS + penicillin-streptomycin was added to each well and explants were incubated overnight at 5% CO_2_ 37°C in a cell culture incubator. Treatments were then added as above, and plates were moved to a Zeiss Cell Discoverer 7 set to an atmosphere of 5% CO_2_ 37°C. Oblique brightfield images were acquired with a 2μm z-step every 30 min for 24hrs with an Axiocam 712 mono camera using the Zen imaging software (version 3.4) and a Zeiss Plan-Apochromat 20x/0.7 autocorr lens. Movies and still images were processed in Zen imaging software.

### Statistical Analysis

Statistical analyses were completed using GraphPad PRISM 9 (Version 9.3.1.) with an α= 0.05 as the threshold for significance. Exact statistical methods used for individual experiments are contained in the figure legends. All graphs represent mean+/- S.E.M. Statistical outliers were determined using a ROUT outlier analyses or Grubbs test in PRISM 9. Reported *n* represent biological replicates (tissue/cells different patient samples) and all experiments were repeated a minimum of three times.

## Acknowledgments

We would like to thank Dr. Carolyn Jones for her feedback on the initial data, Maya Henriquez for the collection of term placental tissue, and all the families that donated tissue for the study. We would also like to thank Dr. Kacie Norton and Arlene Oatway of the University of Alberta Biological Sciences Imaging Facility for their expert technical advice for the electron microscopy sample preparation and imaging. This work was supported by funding from the Canadian Institutes of Health Research (MRC-167968) as well as the Women and Children’s Health Research Institute and their generous donors: the Alberta Women’s Health Foundation and the Stollery Children’s Hospital Foundation. K.P. was supported by the MaTCH program stipend award. S.S is supported by an Alberta-Innovates Graduate Studentship. J.N. was supported by summer studentships from the Women and Children’s Health Research Institute, Alberta-Innovates, and NSERC.

## Competing Interests

The authors have no competing financial and non-financial interests to report.

## Notes

### Competing Interest Statement

The authors have declared no competing interest.

### Summary of Updates

The revised version of the manuscript provides updated results and conclusions.

