## Supplemental Figures for "Loss of polarity regulators initiates gasdermin E mediated pyroptosis in human maternal fetal interface trophoblasts"

#### Supplementary Figure 1:

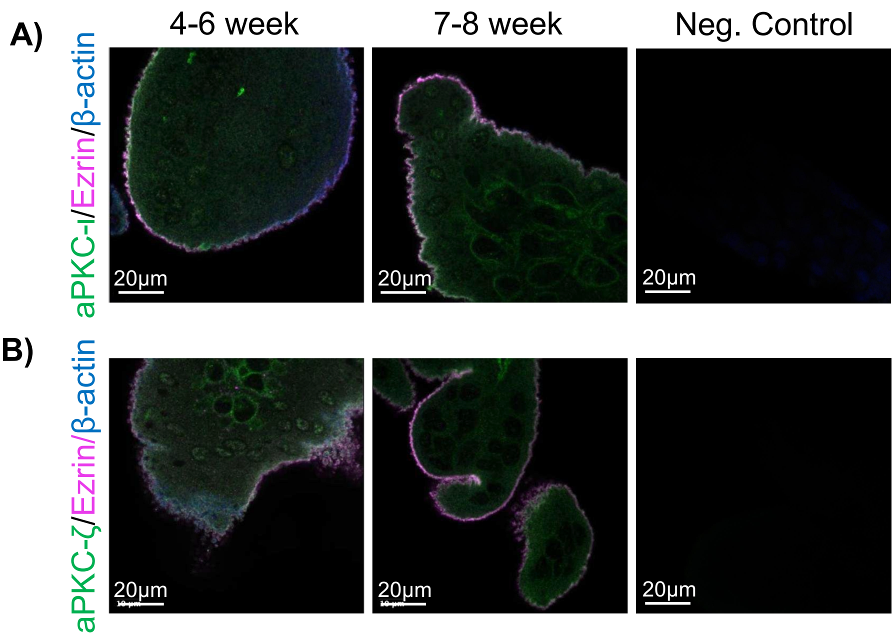


**aPKC-ı and aPKC-ζ colocalize with ezrin apically at 4-6 weeks and 7-8 weeks.** A) Representative images of human placental tissue stained with anti-aPKC-ı (green), anti-ezrin (magenta), and anti-β-actin (blue); left panel=4-6 week; middle panel=7-8 week; right panel=negative control; single plane images from z-stack; B) Representative images of human placental tissue stained with anti-aPKC-ζ (green), anti-ezrin (magenta), and anti-β-actin (blue); left panel=4-6 week; middle panel=7-8 week; right panel=negative control; single plane images from z-stack.

#### Supplementary Figure 2:

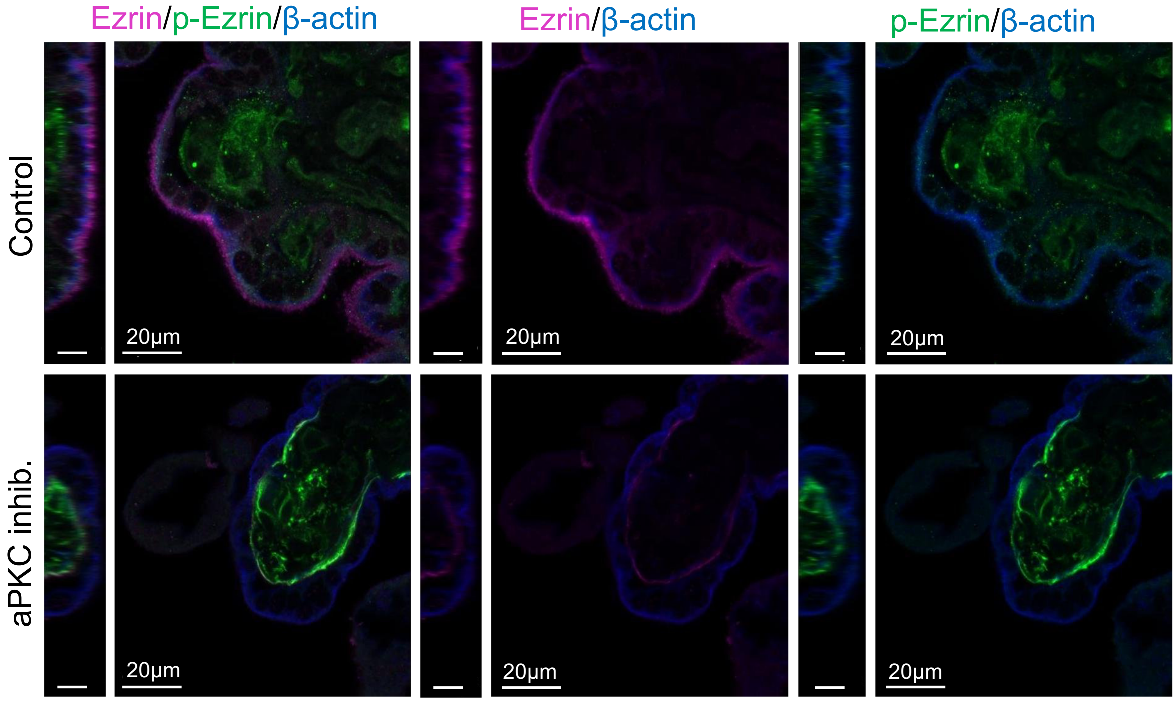


**aPKC inhibitor decreases apical ST ezrin.** Representative images of 37-40 week placental explants treated for 6hrs with aPKC inhibitor and stained with anti-phospho(Thr567)-ezrin (green), anti-ezrin (magenta) and anti-β-actin (blue); left panels=zy plane, scale=10µm; single image plane of z-stack images.

**Supplementary Figure 3:**

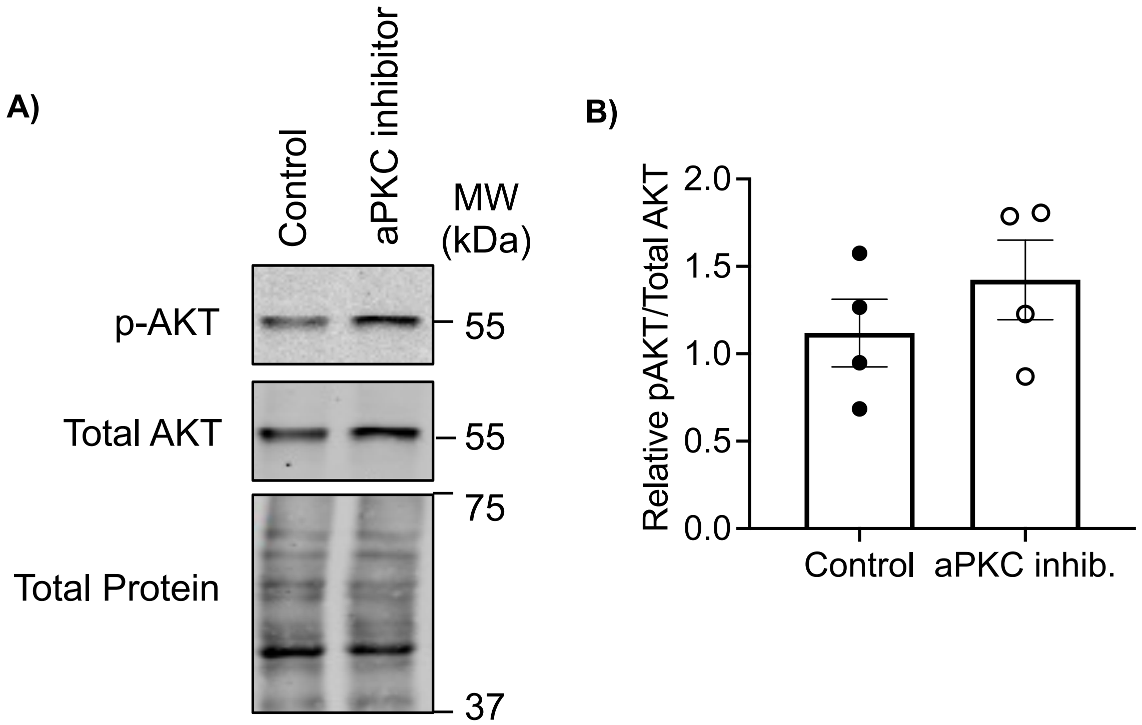


**Treatment of first trimester explants with aPKC inhibitor does not alter AKT activity.** A) Representative western blot with anti-phospho Ser473 AKT, anti-Total AKT, and fast green stain (total protein); B) Summary data of pAKT to total AKT ratios from western blots; *n*=4; Student's t-test; mean +/- S.E.M.

#### Supplementary Figure 4:

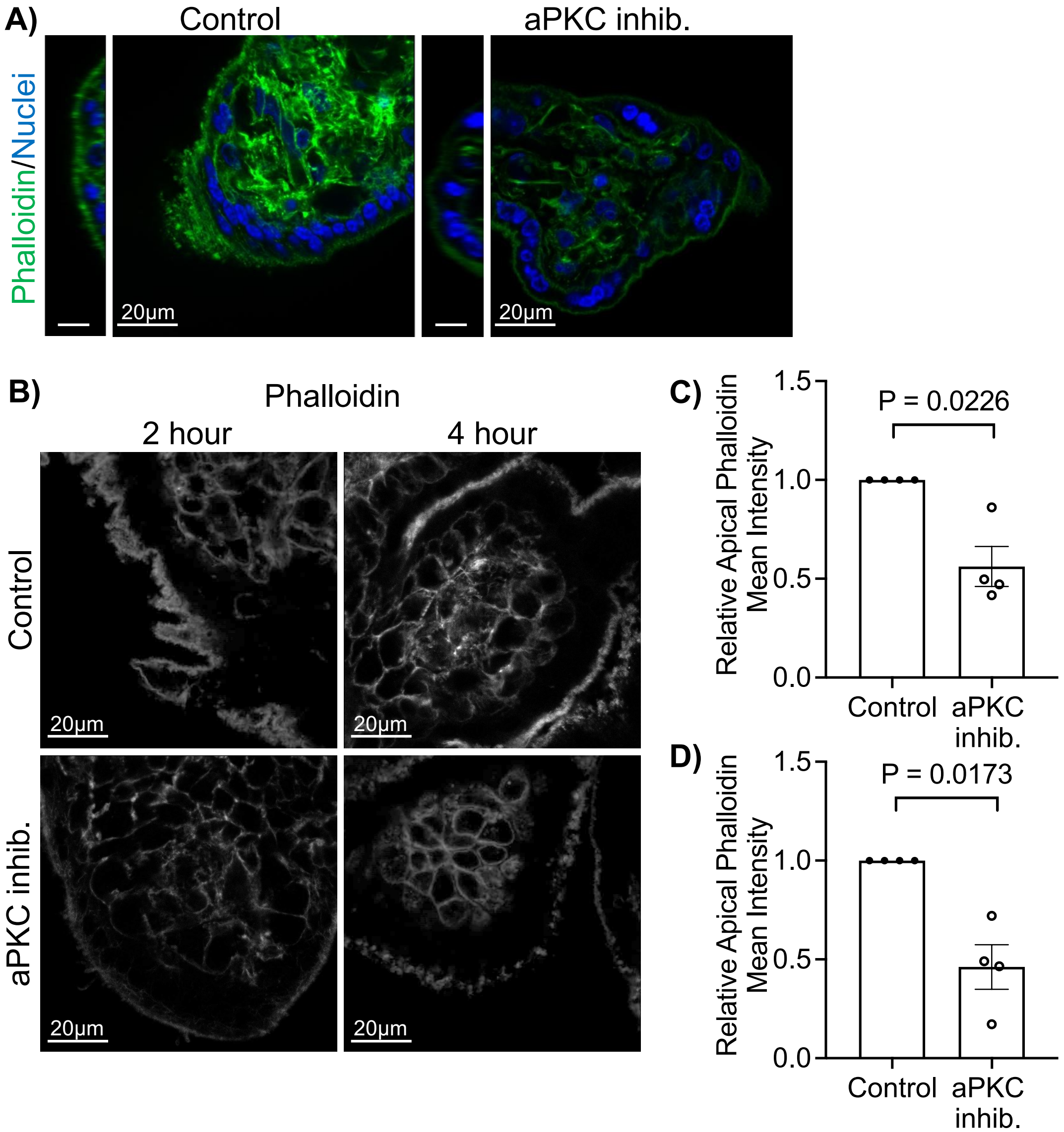


**aPKC inhibitor decreases apical ST phalloidin in first and third trimester placental explants.** A) Representative images of 37-40 week placental explants after staining for phalloidin (green) and Hoescht 33342 (blue) following aPKC inhibitor treatment for 6hrs; yz scale=10µm; single plane image of z-stack; B) Representative images of 9-12 week placental explants after staining for phalloidin (greyscale) following aPKC inhibitor treatment for 2hrs (left panels) and 4hrs (right panels); single plane image of z-stack; C) Summary data for quantitation of apical ST phalloidin mean intensity normalized to control following 2hrs aPKC inhibitor treatment; *n*=4; D) Summary data for quantitation of apical ST phalloidin mean intensity normalized to control following 4hrs aPKC inhibitor treatment; *n*=4; All summary graphs mean +/- S.E.M. with one-sample t-test.

#### Supplementary Figure 5:

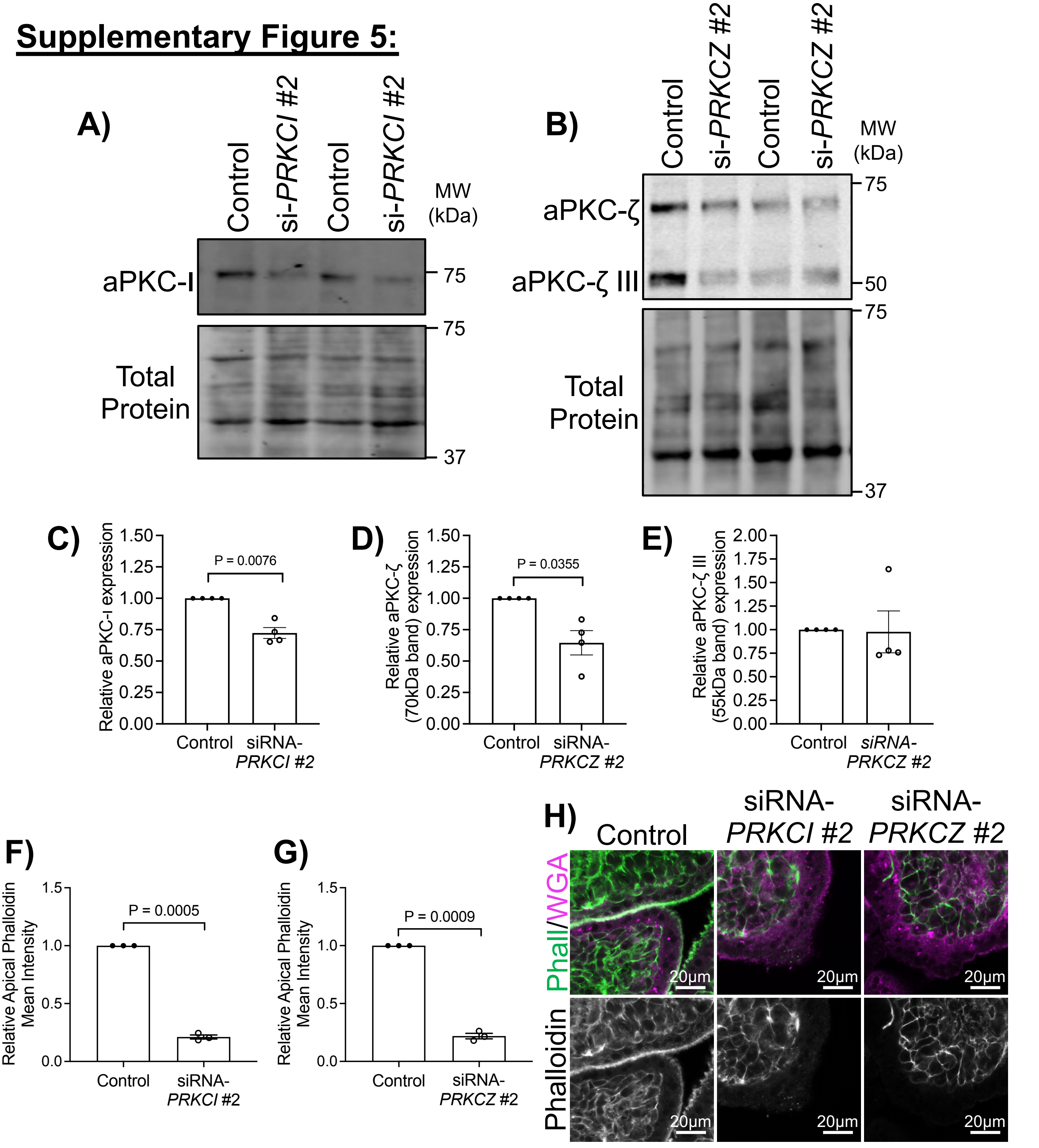


**A second set of siRNAs targeting *PRKCI* and *PRKCZ* significantly decrease ST apical phalloidin intensity.** A) Western blotting of explants treated with *PRKCI* targeting siRNA #2; B) Western blotting of explants treated with *PRKCZ* targeting siRNA #2; C) Summary data for siRNA-*PRKCI* #2 western quantitation*;* D) Summary data for aPKC-ζ quantitation in siRNA-*PRKCI* #2 treated explants; E) Summary data for aPKC-ζ III quantitation in siRNA-*PRKCZ* #2 treated explants; F) Summary data for ST apical phalloidin mean intensity normalized to control in siRNA-*PRKCI* #2 (*n*=3) or G) siRNA-*PRKCZ* #2 treated explants (*n*=3); H) Representative images of phalloidin and WGA (top) or phalloidin (bottom) in siRNA-*PRKCI* #2 or siRNA-*PRKCZ* #2 treated explants; All graphs mean +/- S.E.M. with one-sample t-test.

**Supplementary Figure 6:**

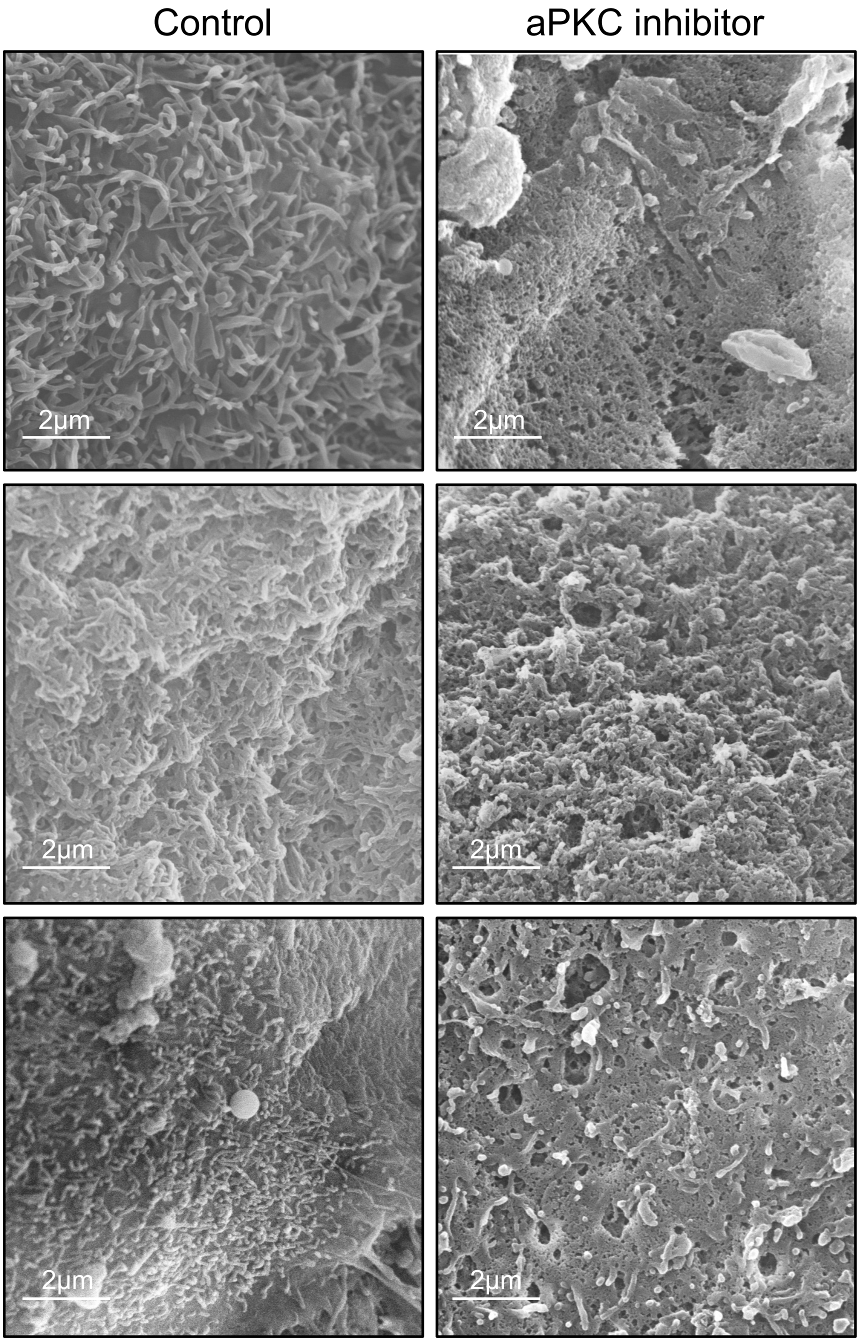


Additional representative SEM images of aPKC inhibitor treated 9-12 week placental explants.

#### Supplementary Figure 7:

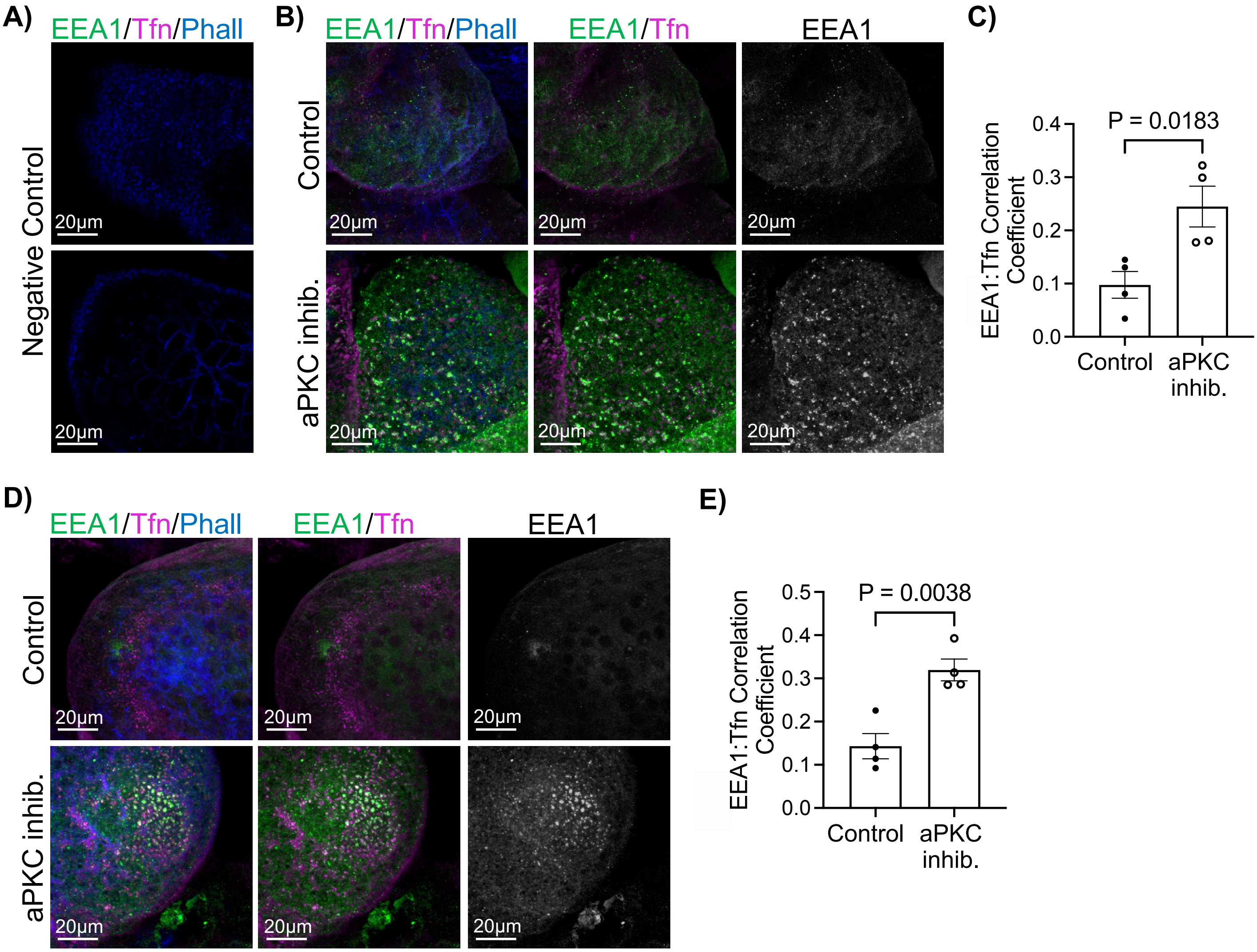


**aPKC inhibitor treatment at 4hrs and 6hrs alters endocytosis of transferrin-594 in placental explants.** A) Representative images of negative control for transferrin and EEA1 staining; phalloidin (blue); B) Representative images of 9-12 week placental explants after transferrin-Texas Red uptake (magenta) and EEA1 (green) staining following treatment with aPKC inhibitor for 4hrs; left panels=merged image; middle panels=EEA1 and transferrin; right panels=isolated EEA1; extended focus images; C) Summary data for quantitation of Global Pearson's coefficient for colocalization between EEA1 and transferrin in 9-12 week placental explants following 4hr aPKC inhibitor treatment; *n*=4; (D) Representative images of 9-12 week placental explants after transferrin-Texas Red uptake (magenta) and EEA1 (green) staining following treatment with aPKC inhibitor for 6hrs; left panels=merged image; middle panels=EEA1 and transferrin; right panels=isolated EEA1; extended focus images; E) Summary data for quantitation of Global Pearson's coefficient for colocalization between EEA1 and transferrin in 9-12 week placental explants following 6hrs aPKC inhibitor treatment; *n*=4; All summary graphs mean +/- SEM; Student's t-test.

#### Supplementary Figure 8:

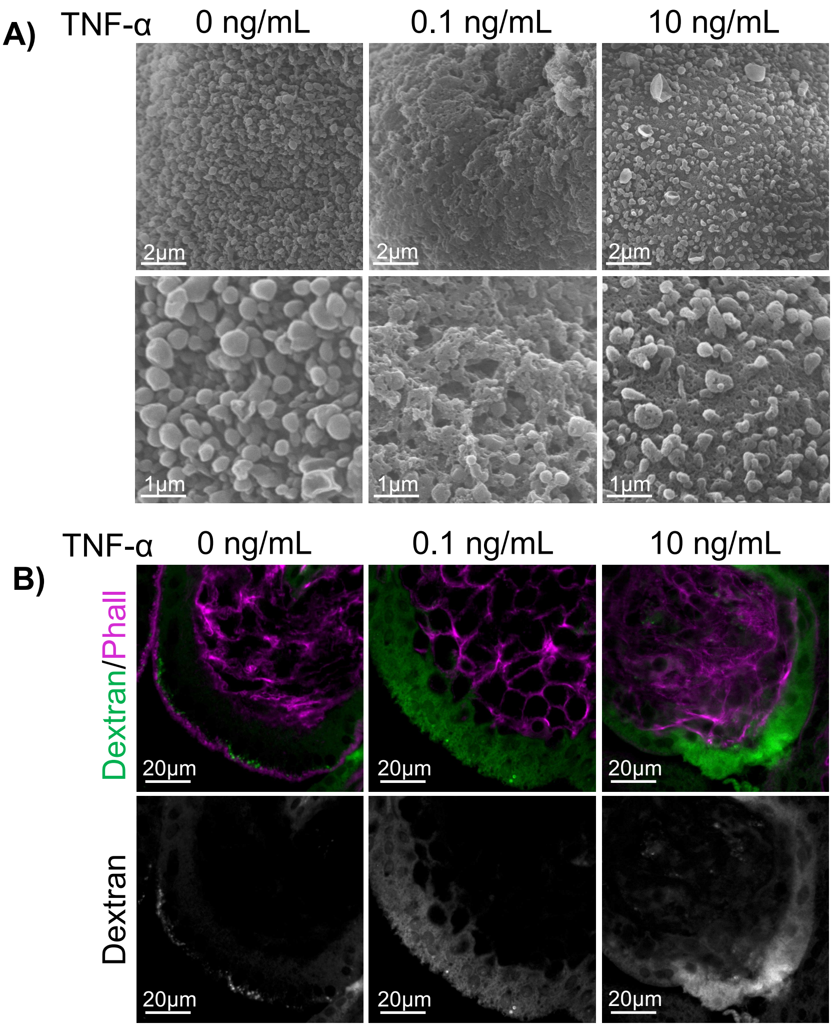


**TNF-α treatment disrupts the ST apical surface structure/morphology and increases ST membrane permeability.** A) Representative SEM images of 37-40 week placental explants following 0-10 ng/mL TNF-α treatment; top panels=lower magnification; bottom panels=higher magnification; B) Representative images of 9-12 week placental explants after incubation with dextran-Texas Red (green) and staining for phalloidin (magenta) following 0-10 ng/mL TNF-α treatment; top panels=merged image; bottom panels=isolated dextran; single plane image of z-stack.

#### Supplementary Figure 9:

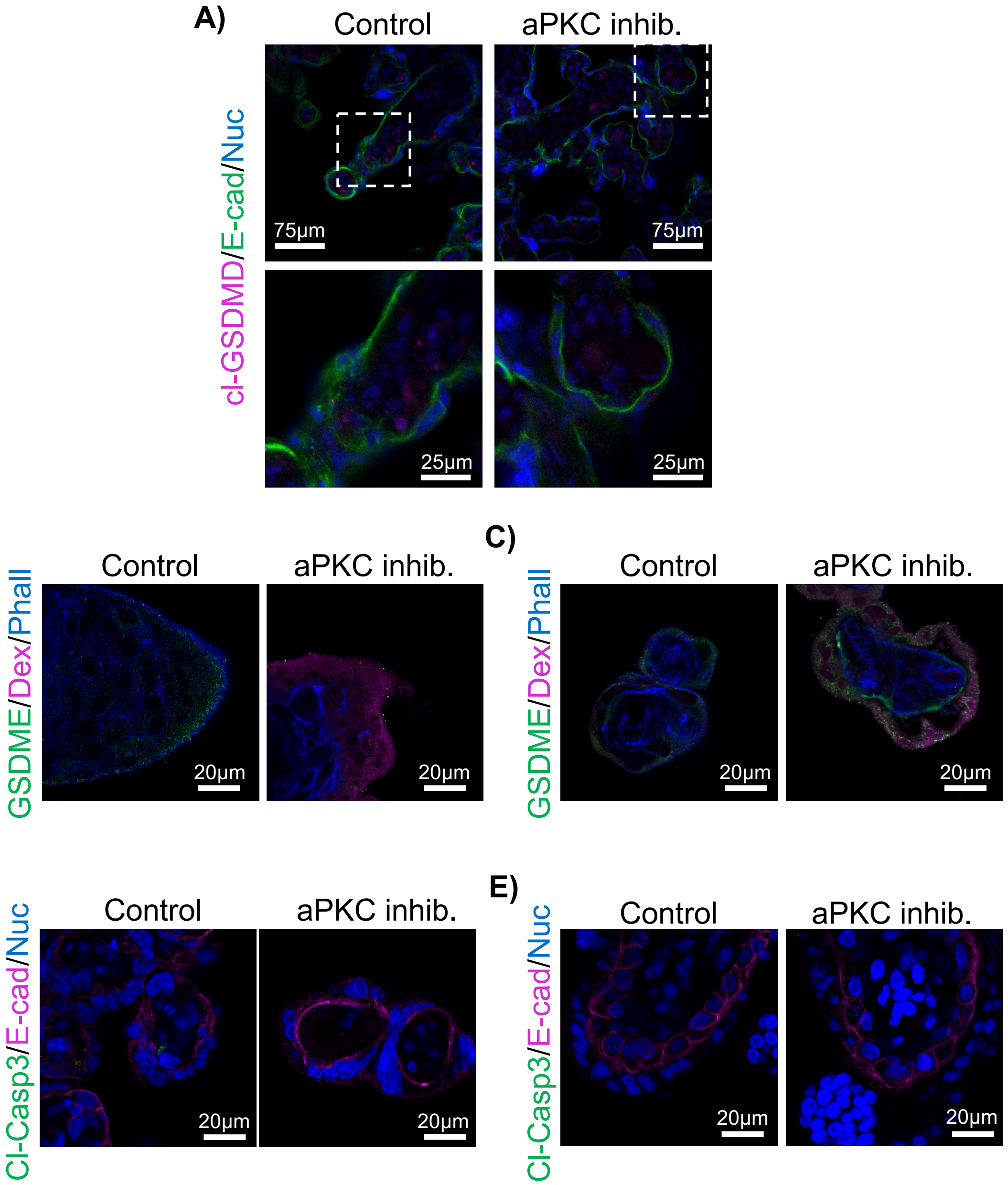


**aPKC inhibitor treatment induces ST pyroptosis.** A) Anti-cleaved gasdermin D (cl-GSDMD; magenta), anti-E-cadherin (green), and Hoescht 33342 in 9-12 week explants treated +/- aPKC inhibitor; bottom panels= higher magnification images of area indicated in top panel; B) Anti-gasdermin E (green), dextran-Texas Red (magenta), and phalloidin; 9-12 week explants +/- aPKC inhibitor treatment; C) Anti-gasdermin E (green), dextran-Texas Red (magenta), and phalloidin; 37-40 week explants +/- aPKC inhibitor treatment; D) Anti-cleaved caspase 3 (cl-Casp3, green), anti-E cadherin (magenta), and Hoescht 33342; 9-12 week explants treated +/- aPKC inhibitor; E) Anti-cleaved caspase 3 (cl-Casp3, green), anti-E cadherin (magenta), and Hoescht 33342; 37-40 week explants treated +/- aPKC inhibitor; All images single xy plane; All treatments 6hrs.

**Supplementary Figure 10:**

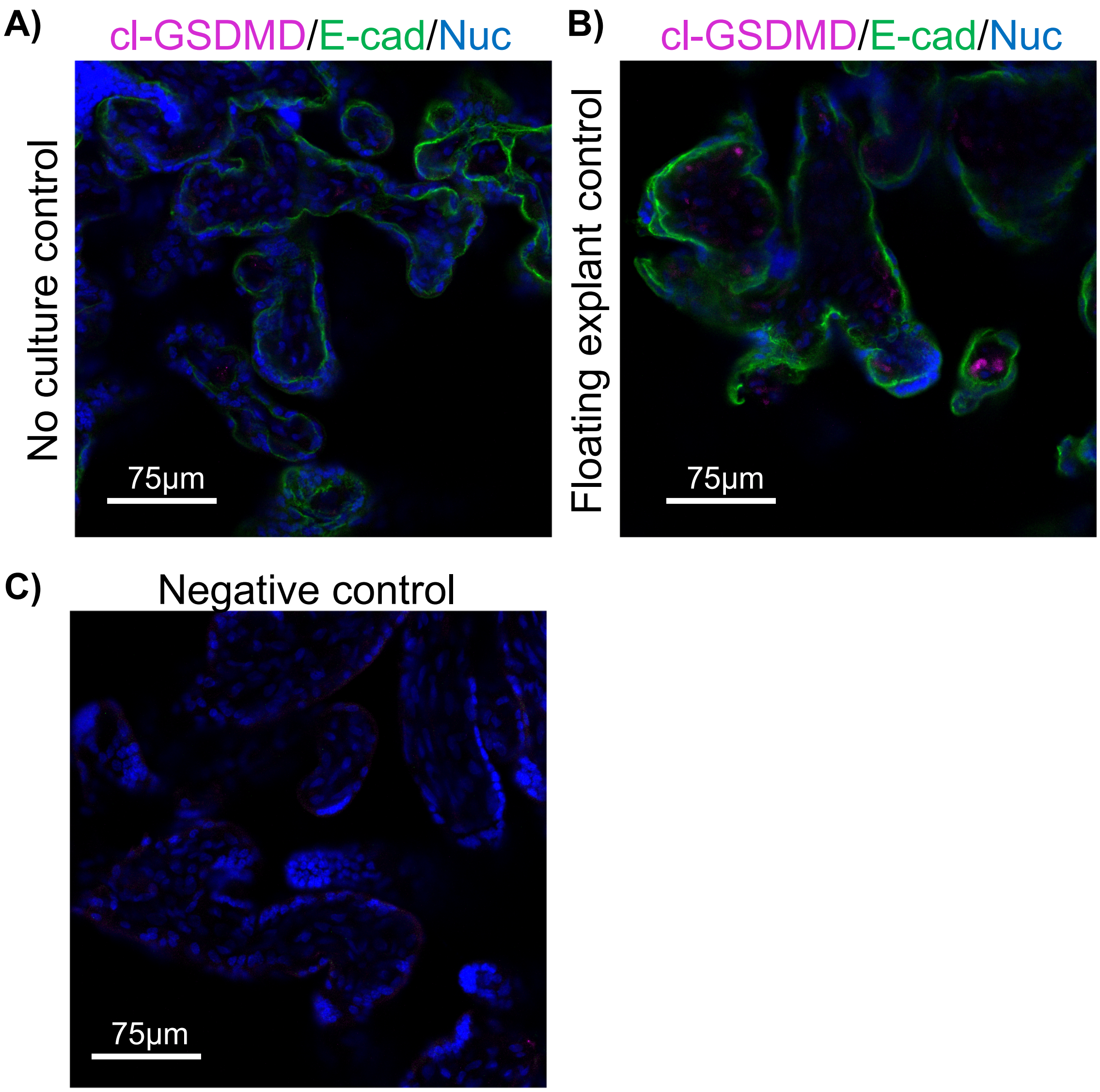


**Gasdermin-D cleavage is observed in the placental stroma of explant cultured tissue but not tissue from the same donor without culture.** A) Term placenta fixed without culture and stained with anti-cleaved gasdermin-D (magenta), anti-E-cadherin (green), and Hoechst 33342 (blue); B) Fixed tissue after floating explant culture from the same donor; C) Negative control.

#### Supplementary Figure 11:

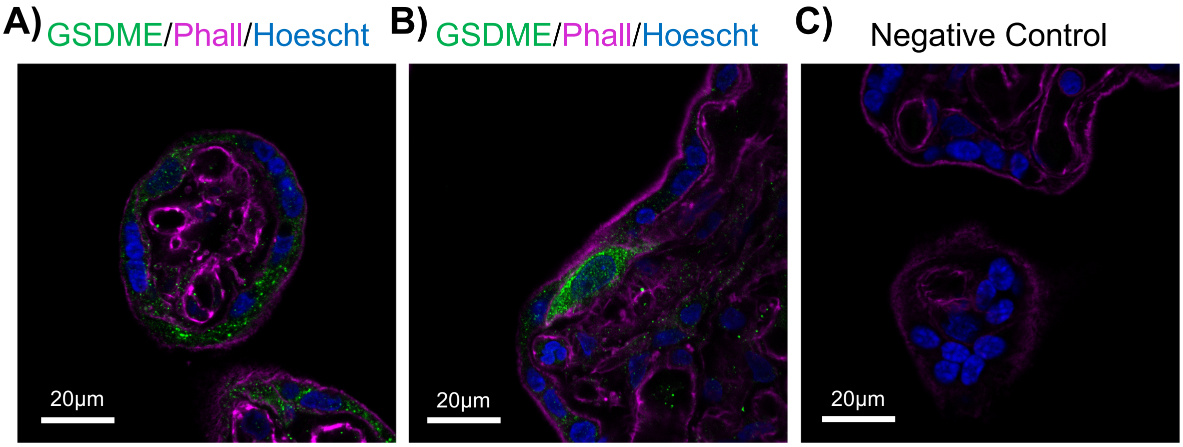


**Gasdermin E is expressed in multiple cell types including the ST in 37-40 week placenta.** A) Terminal villi staining of Gasdermin E (green), Phalloidin (magenta), and nuclear stain Hoechst; B) Strong anti-Gasdermin E signal (green) in a cytotrophoblast progenitor cell; C) Negative control; all images representative of *n=3* patient samples.

<u>**Supplementary Figure 12:**</u>

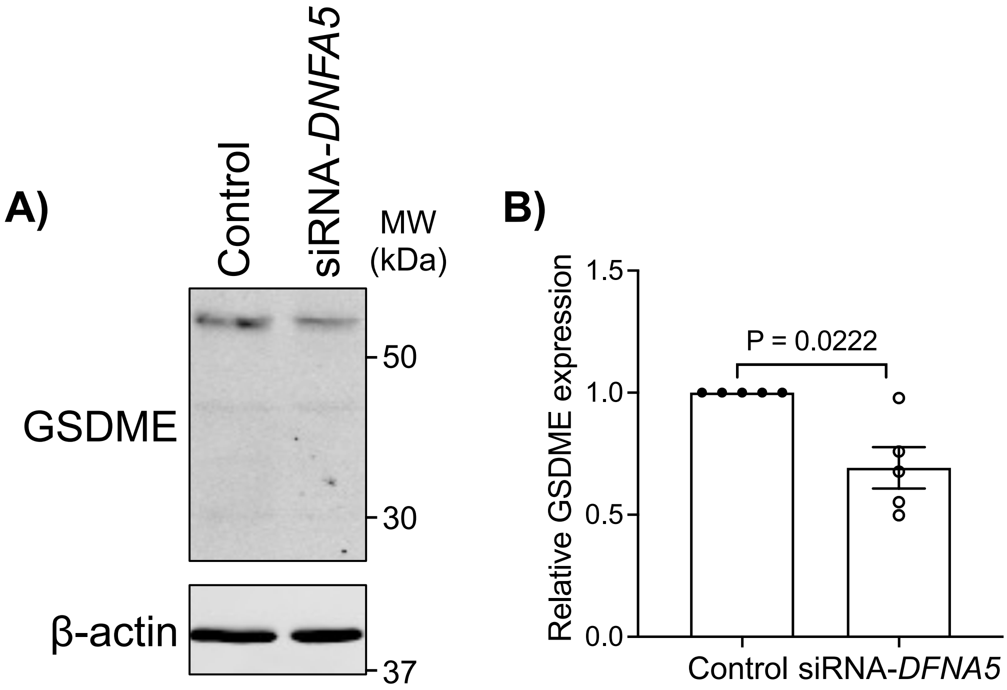


**GSDME targeting siRNA significantly decreases GSDME expression placental explants.** A) Representative western blotting analysis of GSDME targeting (siRNA-*DFNA5*) and negative control treated explant tissue; B) Summary data of GSDME expression quantified by western blotting analyses normalized to control; mean +/- S.E.M.; one sample t-test; *n*=5.

#### Supplementary Figure 13:

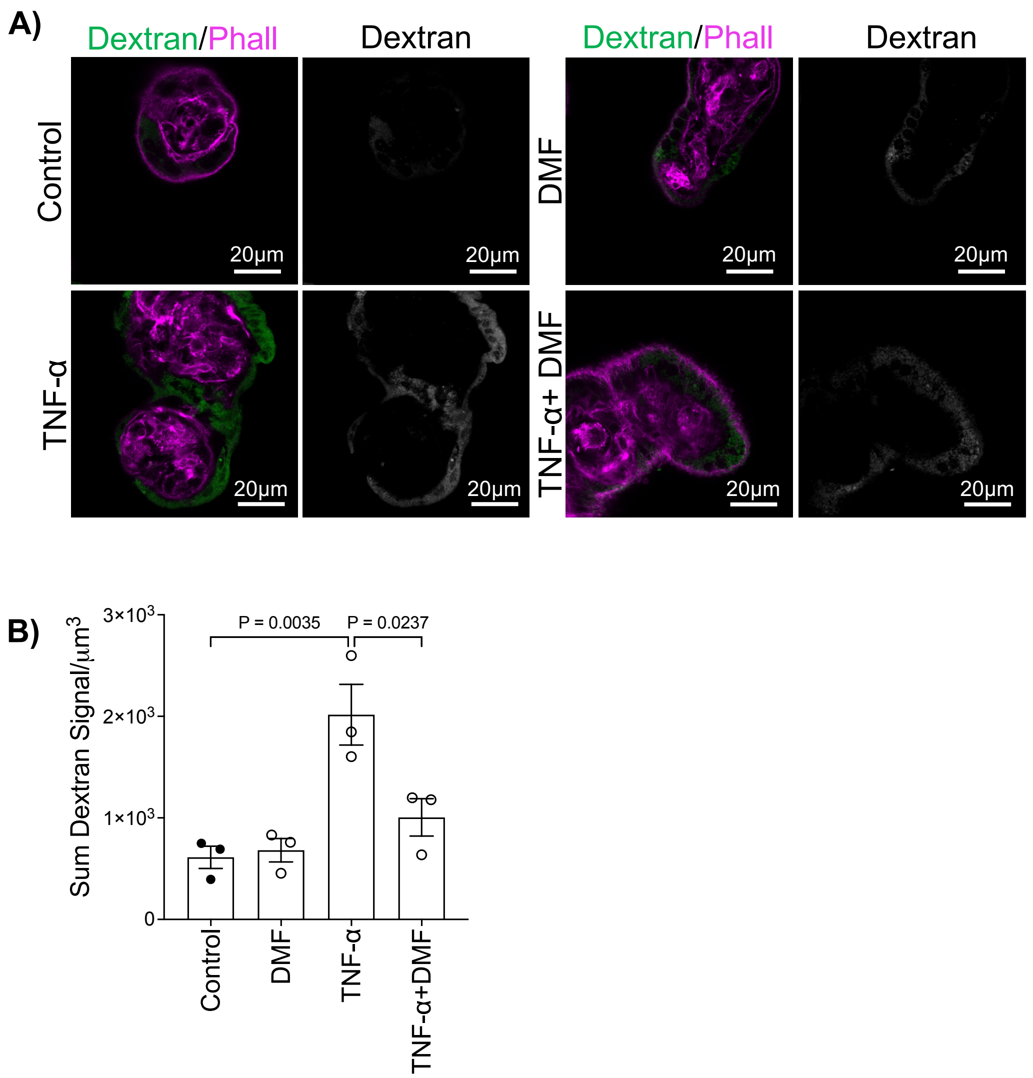


**DMF blocks TNF-α induced ST permeability**. A) Representative images of dextran uptake and phalloidin in 37-40 week placental explants after 6hrs treatment +/- DMF, 100pg/mL TNF-α, or both; right panels=single channel dextran signal; B) Summary data for quantitation of sum dextran signal per $\mu m^3$ in the ST of 37-40 week placental explants; n=3; mean +/- S.E.M.; one-way ANOVA with Sidak's multiple comparison test.

### Source Western blots:

Figure 2I) Anti- aPKC-ι

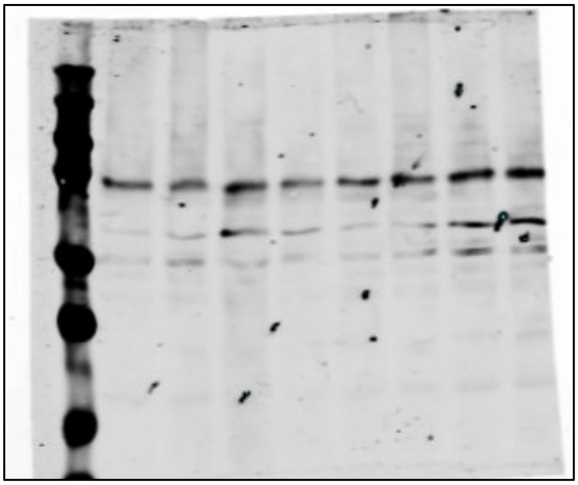

Figure 2J) Anti-aPKC-ζ

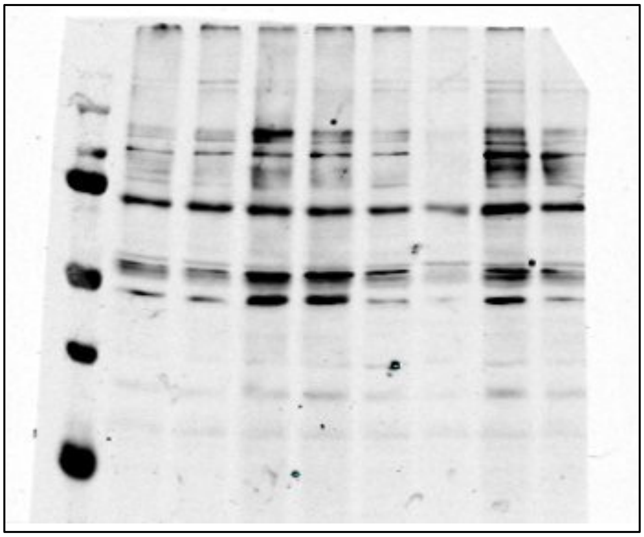

Figure 2K) Anti-Total aPKC

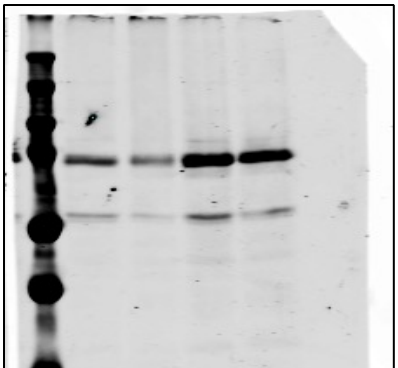

Supplemental Figure 3)

Anti-phospho Ser473 AKT

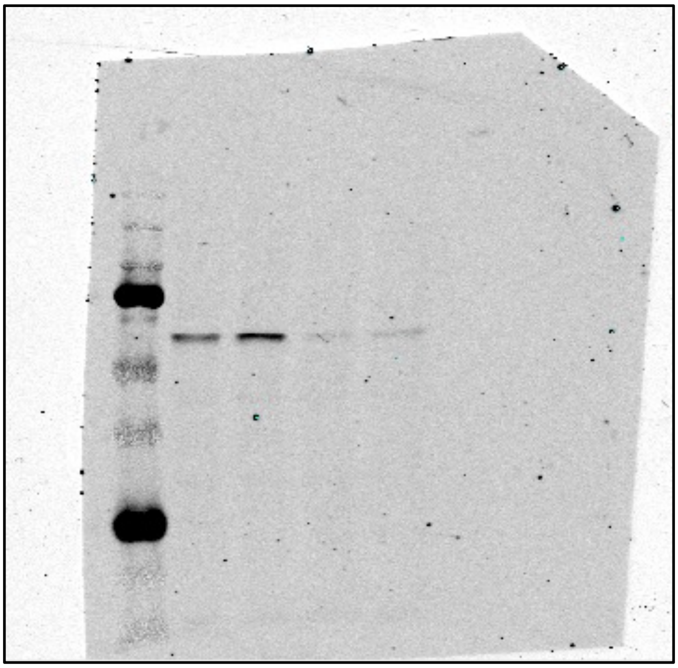

Anti- total AKT

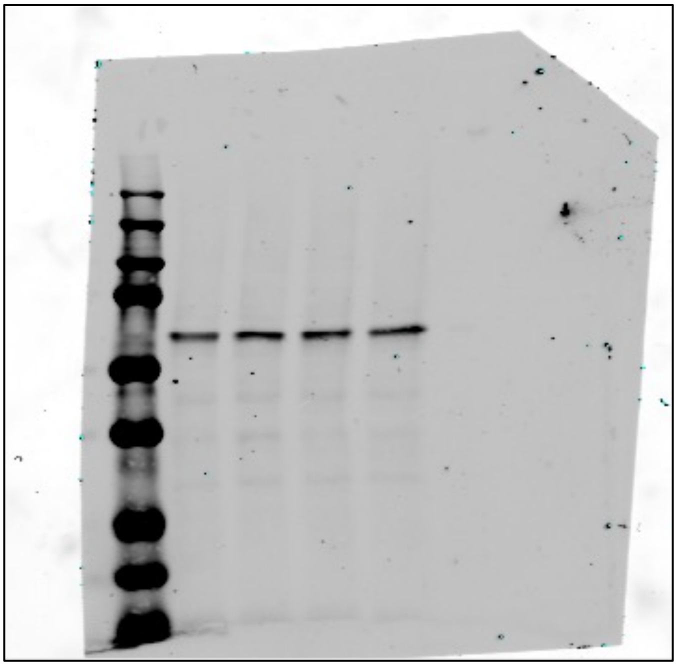

Figure 4A) Anti- aPKC-ι

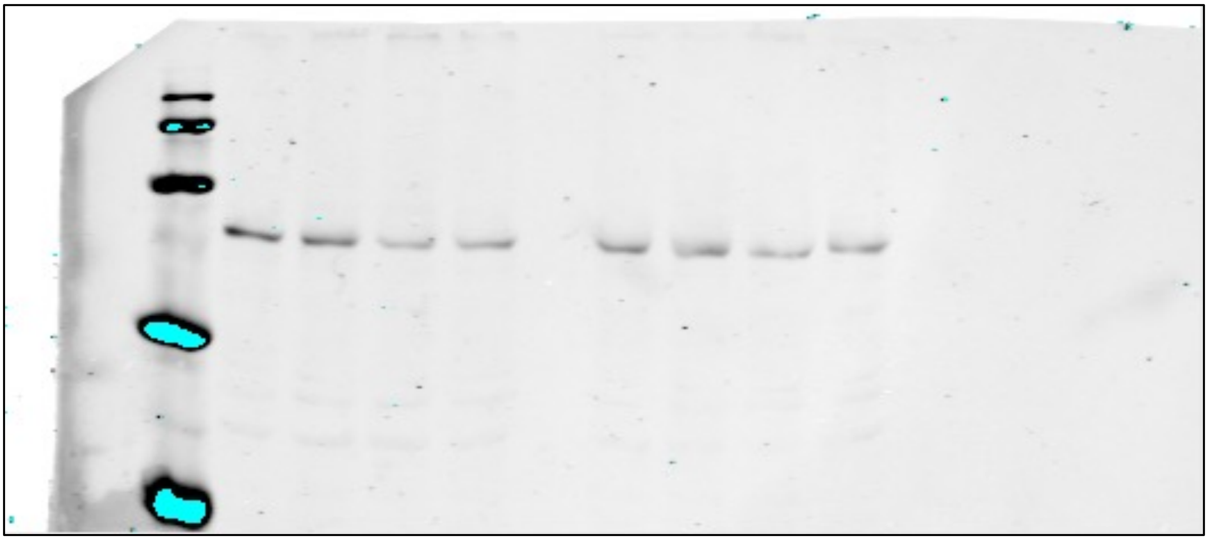

Figure 4A) Anti-aPKC-ζ and anti-β actin

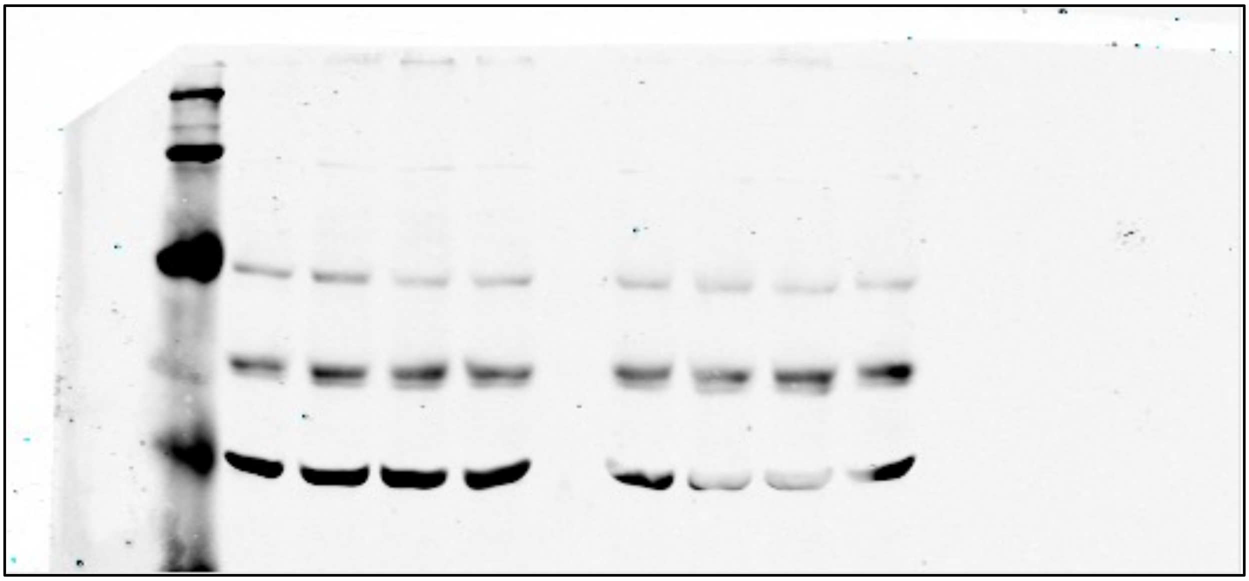

Figure 5G) Anti-GSDME

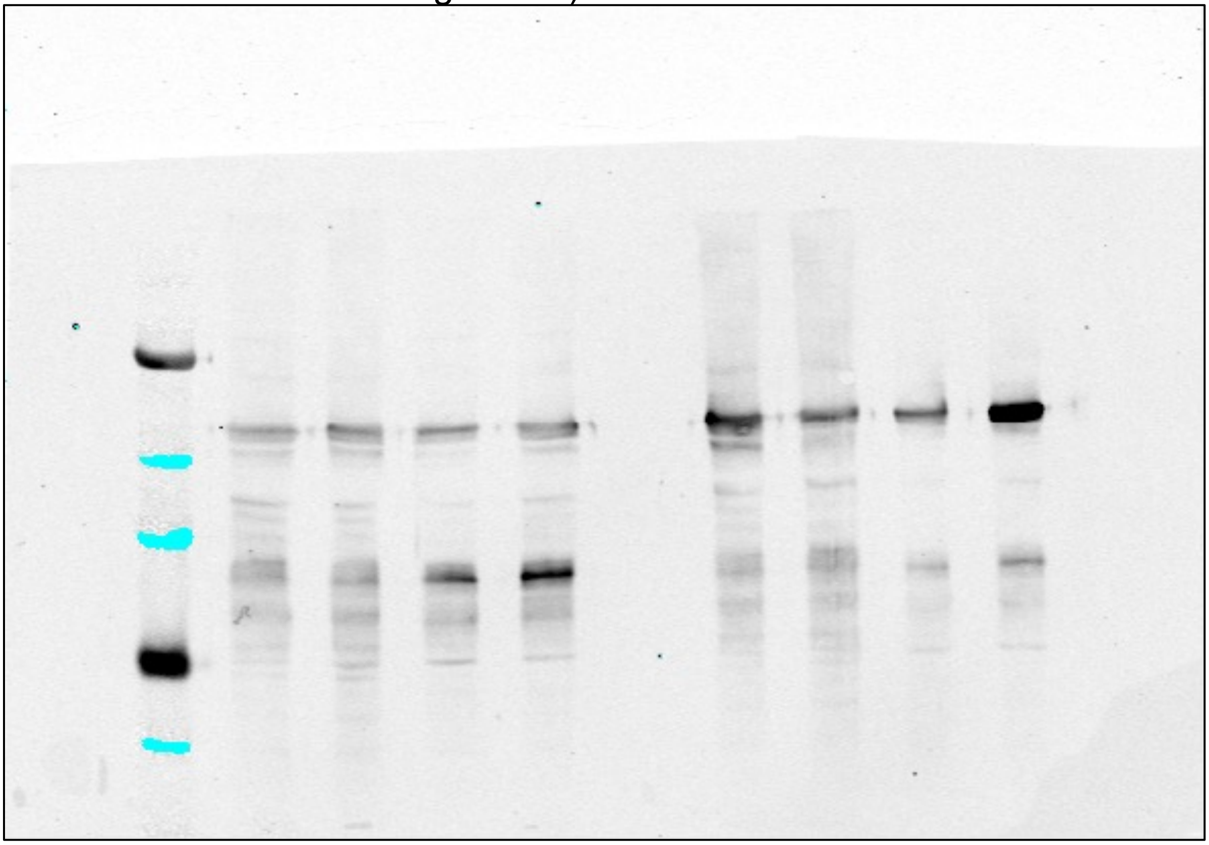

Supplemental Figure 5)

A) Anti- aPKC-ι

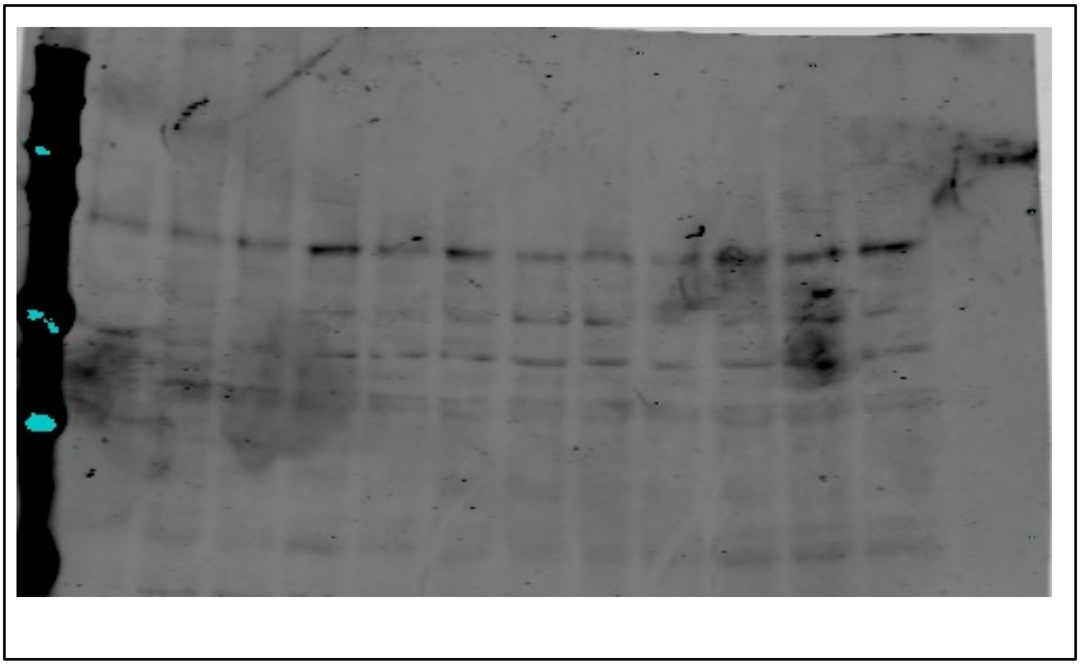

B) Anti-aPKC-ζ

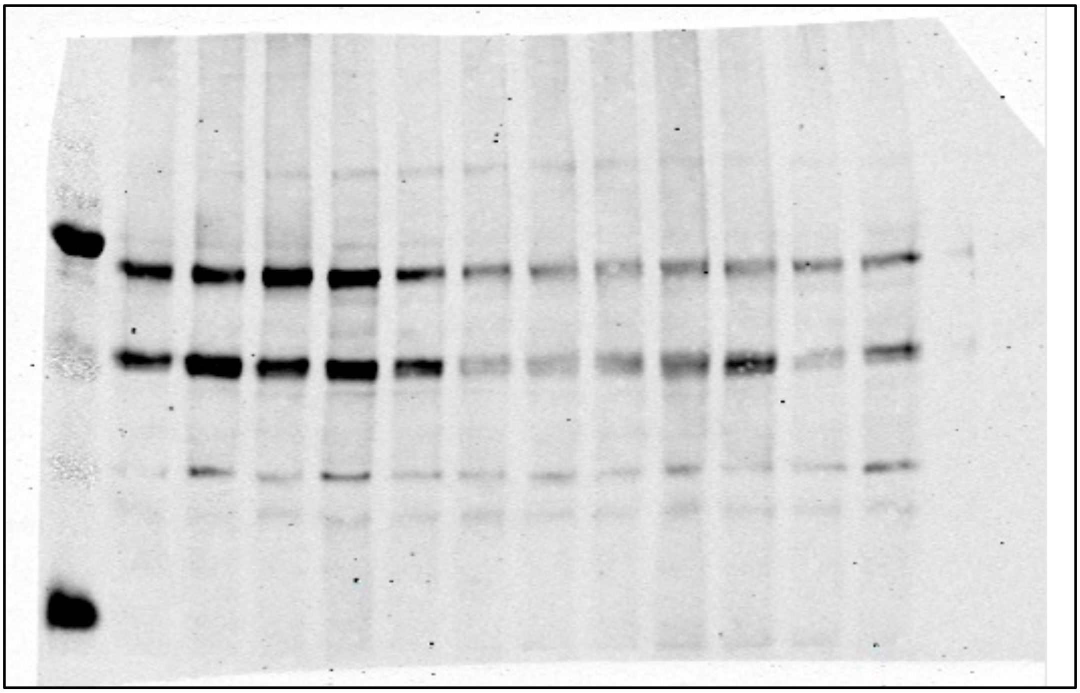

Supplemental Figure 12)

Anti-GSDME

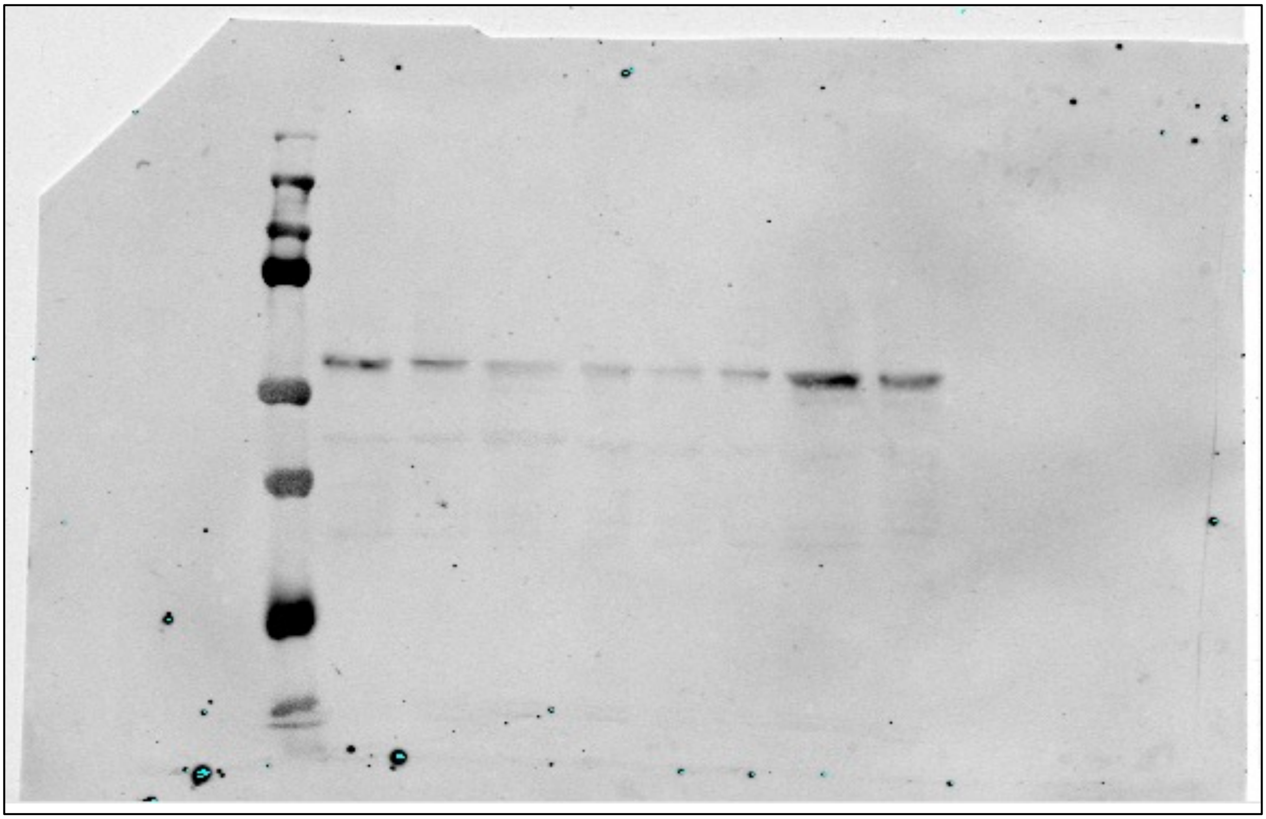
